# EnGens: a computational framework for generation and analysis of representative protein conformational ensembles

**DOI:** 10.1101/2023.04.24.538094

**Authors:** Anja Conev, Mauricio Menagatti Rigo, Didier Devaurs, André Faustino Fonseca, Hussain Kalavadwala, Martiela Vaz de Freitas, Cecilia Clementi, Geancarlo Zanatta, Dinler Amaral Antunes, Lydia Kavraki

**Author notes:** Corresponding authors: *Email addresses:* (Dinler Amaral Antunes), (Lydia Kavraki).

## Abstract

Proteins are dynamic macromolecules that perform vital functions in cells. A protein structure determines its function, but this structure is not static, as proteins change their conformation to achieve various functions. Understanding the conformational landscapes of proteins is essential to understand their mechanism of action. Sets of carefully chosen conformations can summarize such complex landscapes and provide better insights into protein function than single conformations. We refer to these sets as representative conformational ensembles. Recent advances in computational methods have led to an increase in number of available structural datasets spanning conformational landscapes. However, extracting representative conformational ensembles from such datasets is not an easy task and many methods have been developed to tackle it. Our new approach, EnGens (short for ensemble generation), collects these methods into a unified framework for generating and analyzing protein conformational ensembles. In this work we: (1) provide an overview of existing methods and tools for protein structural ensemble generation and analysis; (2) unify existing approaches in an open-source Python package, and a portable Docker image, providing interactive visualizations within a Jupyter Notebook pipeline; (3) test our pipeline on a few canonical examples found in the literature. Representative ensembles produced by EnGens can be used for many downstream tasks such as protein-ligand ensemble docking, Markov state modeling of protein dynamics and analysis of the effect of single-point mutations.

## 1. Introduction

Proteins are the main building blocks of cells, executing a variety of functions vital to life, such as signal transduction, immune defense, and DNA replication. These functions are driven by the three-dimensional arrangement (i.e., the structural conformation) of proteins [1]. However, proteins exist in a highly complex environment and are not static entities. The following examples demonstrate that a single protein conformation is not enough to characterize important protein dynamics driving diverse functions. First, allosteric modulations, driven by mutations or drug interactions far from the protein’s active site, induce conformational changes within the active site [2], which can modify the protein’s activity [3–5]. Second, metamorphic proteins [6–8] switch between drastically different folds of the same sequence, thereby performing different functions [9]. Finally, intrinsically disordered proteins and intrinsically disordered protein regions constitute extreme examples of highly flexible structures. They exist as highly dynamic structural ensembles [10] failing to form a globally stable three-dimensional shape in physiological solution, thereby performing different functions. All these examples demonstrate the importance of comprehensively characterizing a protein conformational landscape and identifying key conformational states to understand protein function [11].

The energy landscape theory [12–14] is one framework that provides an understanding of protein structure and dynamics by analyzing a protein’s free energy landscape (or free energy surface - FES) as a function of a few collective variables (CVs). However, the exact determination of the FES for large proteins is challenging, as it requires extensive sampling of the protein’s conformational space. New methods for computational protein structure prediction [15–17] and simulation [18– 22] are emerging and there is an increased availability protein structure datasets. However, a full understanding of a protein’s dynamics can be reached only when the dataset spans the FES sufficiently, allowing quantitative methods (such as Markov state modeling) to be applied. In this work, we do not tackle the sampling problem, as we rely on previously generated datasets. In other words, our approach focuses on structurally representative ensembles and not on thermodynamic ensembles.

There is a need to rapidly extract useful information from conformational datasets [23] without directly modeling the dynamics. Subsets of conformations extracted to represent major conformational states contained within the data provide a useful conformational summary. We call such sets *representative conformational ensembles*. Note that, in this context, the term ensemble does not refer to a statistical ensemble. The task we address is that of extracting described representative conformational ensembles from datasets of protein structures. Extracted representative ensembles can be useful for many downstream tasks such as protein-ligand ensemble docking [24], analysis of mutational effects [25] and extensive Markov state modeling of protein dynamics [26, 27]. It is important to provide sufficient analysis of the extracted ensemble to summarize important properties of each protein state (e.g., the distance between protein domains or the distance between important residues in the active site) and help derive more intuitive insights (e.g., whether a member of the ensemble represents the protein in its active or inactive form). In this work, we develop EnGens - a computational pipeline for the generation and analysis of representative protein conformational ensembles.

Sources of protein structural datasets are now diverse. The Protein Data Bank (PDB) [28, 29], first established in 1971, has experienced steady growth over the past decade. With more than 10,000 experimentally solved protein structures deposited annually, the total number of available entries to date is around 200,000. These data have allowed for new breakthroughs in the field of protein structure prediction, including machine learning techniques such as AlphaFold2 [15], RosettaFold [16] or ESMFold [17]. The AlphaFold database [30] was released with over 200 million protein structure predictions. ESMFold has recently reported comparable performance to AlphaFold2 with the ESM Metagenomic Atlas, which contains 617 million predicted metagenomic structures. This vast amount of available data allows researchers to collect multiple conformations of the same protein [31]. Collected conformations make up datasets whose content can be summarized and analyzed with EnGens. We call such datasets “static” to highlight the fact that the conformations they contain are independent and not derived from simulating protein dynamics.

A more extensive analysis of protein dynamics can be performed using simulations that generate so-called “dynamic” datasets. Conformations within these datasets are not independent - they are time-ordered and form trajectories. Molecular dynamics (MD) simulations, first developed in the late 70s [32], have been established as a gold standard for exploring protein dynamics. Many computational packages have since been developed, including NAMD [33], GROMACS [34, 35], AMBER [36], CHARMM [37], OpenMM [38]. MD software is becoming more accessible with python plugins and graphical user interfaces [18]. Markov State Modelling (MSM) approaches for interpreting MD simulations [39] have recently gained popularity. but constructing MSMs can be a lengthy process as this requires extensive sampling. On the other hand, EnGens can be used to gain insights into the content of MD datasets without fully modeling the dynamics.

Our approach recognizes and addresses the need for a unified computational framework to help researchers summarize the vast amount of newly available structural data in an effort to understand the conformational landscape driving protein function. EnGens builds on several existing tools that have proven useful for protein structure analysis. For the computational representation of protein structure, EnGens utilizes the PDB module of BioPython [40] as well as the rich featurization module of PyEmma [41], powered by MDTraj [42]. For dimensionality reduction and clustering steps EnGens provides a diverse set of algorithms implemented across deeptime [43], scikit-learn [44], UMAP [45] and SRV [46]. EnGens brings all these tools closer to the community by providing an open-source pipeline wrapped into a portable Docker image and accompanied by extensive example workflows written in Jupyter Notebooks. Additionally, EnGens implements a set of customizable interactive visualizations that provide users with detailed insight into the generated conformational ensembles.

Other similar tools complement EnGens (Table S1). CoNSEnsX [47] generates ensembles based on available NMR data. PENSA [48, 49] provides different metrics (Jensen-Shannon Distance, Kolmogorov-Smirnov Statistic, Overall Ensemble Similarity) for the comparison of generated ensembles. ProDy [50, 51] provides a set of algorithms for studying protein dynamics, which includes normal mode analysis. The specificity of EnGens lies in that: (1) it provides customizable PyEmma featurization for both static and dynamic datasets; (2) it contains both linear and nonlinear dimensionality reduction techniques (linear PCA [52] and TICA [53, 54]; nonlinear UMAP and SRV); (3) it provides different clustering methods (hierarchical, K-means and Gaussian Mixture Models); (4) it is wrapped in an accessible Docker image and includes interactive Jupyter Notebook Workflows with rich ensemble visualizations. With these unique properties, EnGens enables users to automate the generation and analysis of protein conformational ensembles. We envision EnGens as an important and useful resource for data analysis of protein structure to support researchers in the era of big data.

In the following sections, we describe methods involved in the EnGens pipeline. Note that these methods have been previously published and extensively validated [41, 45, 46, 53, 54]. Hence, validating these methods is outside of the scope of our work. Instead, we showcase the use of the full EnGens pipeline on a set of example molecules from the literature for which static or dynamic datasets are available. This includes molecules of different scales: a large protein complex (PI3K kinase), a peptide drug (Compstatin) and a small molecule (Nelfinavir).

## 2. Methods

We have developed EnGens, an automated pipeline for generating and analyzing protein conformational ensembles, given a dataset of protein structures as input. Note that EnGens pipeline has two distinct use-cases: i) processing static protein datasets (e.g., experimental structures); ii) processing dynamic protein datasets (e.g., MD simulations).

A static structural dataset could be experimentally derived and collected from the PDB or modeled computationally (e.g., using AlphaFold or Modeller). For a dataset extracted from the PDB, EnGens can be used to reveal different conformational states and extract a representative ensemble summarizing the dataset. For a dataset compiled by computationally modeling a protein and its common mutants, EnGens can describe the conformational landscape of mutants and help point out the impact of mutations.

A dynamic structural dataset is generally a trajectory derived from an MD simulation. If the simulation involves a protein with a ligand in its active site, EnGens can point out conformational changes that occur upon binding. It is important to note that for the analysis of MD-derived data much work has been done in the field of Markov state modeling [26, 55, 56]. Modeling the dynamics of a system is outside of the scope of EnGens pipeline as its goal is only to generate and analyze the representative conformational ensemble. However, the dynamic use-case is largely inspired by the insights from Markov modeling approaches. For example, one important insight is that resolving slow processes can help identify biologically relevant conformational changes. Thus, using methods related to Markov modeling helps EnGens uncover conformational states and ensembles with biological relevance.

Both static and dynamic datasets can potentially include large numbers of structures that are difficult to systematically inspect visually. To address this issue, EnGens partitions the structural dataset into clusters and extracts a representative conformation from each cluster to form a structurally diverse conformational ensemble. The EnGens pipeline is divided into four workflows that are summarized in Figure 1. Below we give an overview of the workflows and their respective goals. A detailed description of each workflow is provided in the supplementary text.

**Figure 1:**
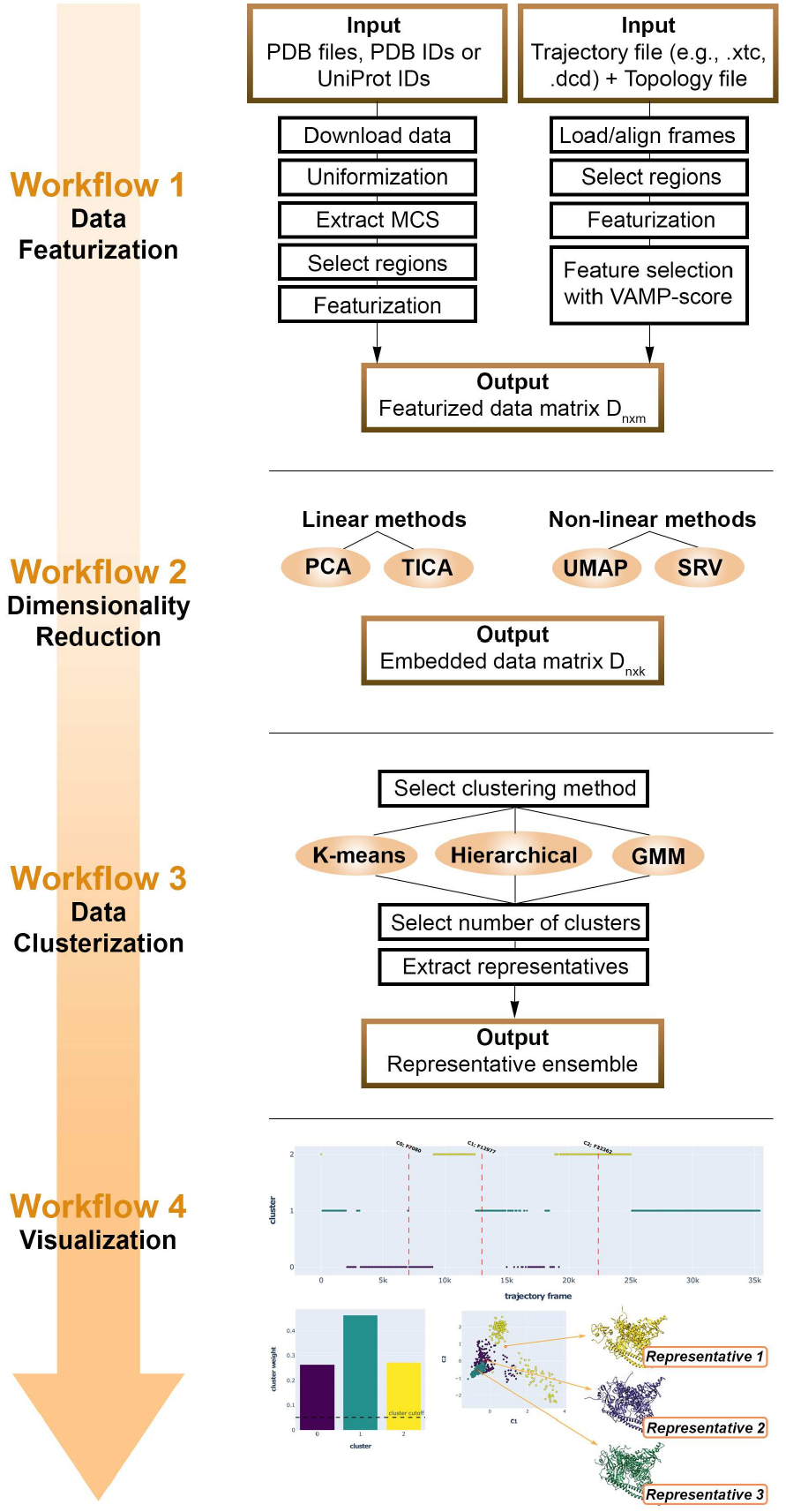
Overview of the EnGens methodology. Workflows are listed across the vertical arrow on the left. Individual steps are listed in the diagram on the right.

### 2.1. Workflow 2: Projecting the featurized representation into an embedding in low dimensional space

Numerical vectors extracted from the first workflow often have very high dimensionality. Depending on the size of the protein and the type of featurization, this vector could contain thousands of elements to represent one structure. High dimensional data presents unique challenges for clustering algorithms, as metrics lose their utility in high dimensional spaces. It is thus important to embed the information into a lower dimensional space before clustering. In Workflow2 we provide implementations of four widely used algorithms for dimensionality reduction. For the static use-case we provide two standard methods: principal components analysis (PCA) and uniform manifold approximation and projection (UMAP). For the dynamic use-case we provide two additional methods that make use of the time ordered nature of the data: time-lagged independent components analysis (TICA) and state-free reversible VAMPnet (SRV). TICA and SRV are not suitable for static datasets, which lack the time component that is exploited by these methods. TICA and PCA are linear methods, while UMAP and SRV are non linear and can thus identify non linear relationships between features. The result of Workflow2 is an embedding of the data in a lower dimensional space, in which the data can be more efficiently partitioned into clusters to identify a representative ensemble.

### 2.2. Workflow 3: Clustering embeddings and extracting the ensemble

Low dimensional embeddings represent each conformation in the dataset. Various distance metrics can be used to calculate similarity between two conformations. This allows us to identify clusters of similar datapoints. In Workflow 3 we provide implementations of three widely used clustering algorithms: hierarchical clustering [57], K-means [58] and gaussian mixture models (GMM) [59]. Hierarchical clustering provides a dendrogram of the data, allowing users to visually inspect the clusters and their relationships. The lower computational complexity of K-means makes it more suitable for large datasets. While K-means assumes a spherical data distribution, GMM can handle more complex distributions and provide a probabilistic model. Whatever the method, resulting clusters correspond to groups of structurally similar conformations. Then, we find the hub of each cluster as the point with the most neighbors and call it a cluster representative. Finally, we generate the resulting ensemble by extracting cluster representatives.

### 2.3. Workflow 4: Visualizing the data and analyzing the ensemble

In the final workflow we provide a set of customizable interactive plots to analyze the generated ensemble. Users can visualize and inspect the 2D embeddings and their clustering. The extracted representatives are highlighted and their position within the 2D embedding space can be identified. Additionally, users can visualize the 3D atomic-resolution conformations of the extracted representatives. The ensemble can be further analyzed by generating a scatterplot of interesting features (e.g., the distance between important residues or RMSD to a template conformation) for each conformation. The same information can be summarized per cluster as a box plot. These visualizations are meant to help users interpret the ensemble (e.g., understand if the active and inactive states of a protein are represented within the ensemble).

## 3. Results

The algorithms gathered under the umbrella of the EnGens pipeline have been validated in past literature [41, 45, 46, 53, 54]. The validation of these methods being therefore outside of the scope of this work, in this section we showcase the use of the full EnGens pipeline. To this end, we have selected proteins for which structural data had been analyzed manually via often laborious processes to extract a conformational ensemble. We process the data entirely within the EnGens pipeline, and show that we can generate the same conformational ensemble as reported in previous studies. The examples we picked cover three systems of varying complexities. First, we process a large PI3K protein complex within both use-cases: a crystal structure dataset and an MD trajectory. Second, we apply EnGens to an MD trajectory of the peptide ligand Compstatin. Finally, we use the same methodology to process an MD trajectory of the small drug Nelfinavir.

### 3.1. Class I PI3K (PI3K-I) experiments

PI3K-I is a family of lipid kinase proteins that phosphorylate a lipid found on the plasma membrane, regulating cell growth and proliferation [60]. Increased activity of PI3K-I has been associated with oncogenesis and its the structural aspects have been widely studied. Members of PI3K-IA subfamily contain a regulatory (p85) and a catalytic (p110) subunit (Figure S1). Kinase activity is autoinhibited by the interaction between the nSH2 domain of the regulatory unit and the C2 domain of the catalytic unit [61, 62]. For instance, it has been shown that the nSH2 domain moves away from the catalytic unit upon contact with a phosphorylated tyrosine pY of the receptor tyrosine kinase (RTK). This movement leads to the activation of PI3K-IA [63–65]. Two recent works performed further structural analysis of the PI3K, one using available PI3K crystal structure data [66] and another performing and analyzing MD simulations of a mutant [67].

#### 3.1.1. PI3K-IA: crystal structure dataset

We base this experiment on a study by [66] that extracted from the PDB a dataset of 49 dimer structures corresponding to alpha, beta and delta isoforms of PI3K-IA (Table S4). While all structures are dimers (containing both catalytic and regulatory units), they differ in the portion of the regulatory unit that is crystalized, namely the nSH2, iSH2 and cSH2 domains. 10 structures were crystalized without the nSH2 domain (PI3KΔnSH2), while the rest contain the nSH2 domain (PI3K+nSH2). The analysis by Zhang et al. was performed by manually engineering the feature of interest as distance between the C2 domain and the kinase domain of the PI3K catalytic unit. With a manually set threshold they divide the 49 structures in two groups: active/open (12) and inactive/closed (37). Zhang et al. conclude that all 10 PI3KΔnSH2 structures have nSH2 released and are active/open. Additionally, two of the PI3K+nSH2 structures have a mutation that leads to the activation. The other 37 structures are considered autoinhibited and are labeled inactive/closed. When processing this dataset with the EnGens pipeline, our goal was to test the ability of EnGens to generate a diverse ensemble of structures that would include representative structures of the active and inactive states. We use PDB codes of the dataset as input (Table S4). EnGens extracts the maximum common substructure (MCS) for each structure (see Supplementary Material: Workflow 1S-2). The MCS includes the catalytic unit and the iSH2 domain of the regulatory unit (Figures S2, S3, S4). We featurize each structure by using the pairwise distances between the centers of mass of the MCS chains. We choose the PCA option for dimensionality reduction step and K-means for clustering. Results of the analysis as provided by the EnGens dashboard are shown in Figure 2.

**Figure 2:**
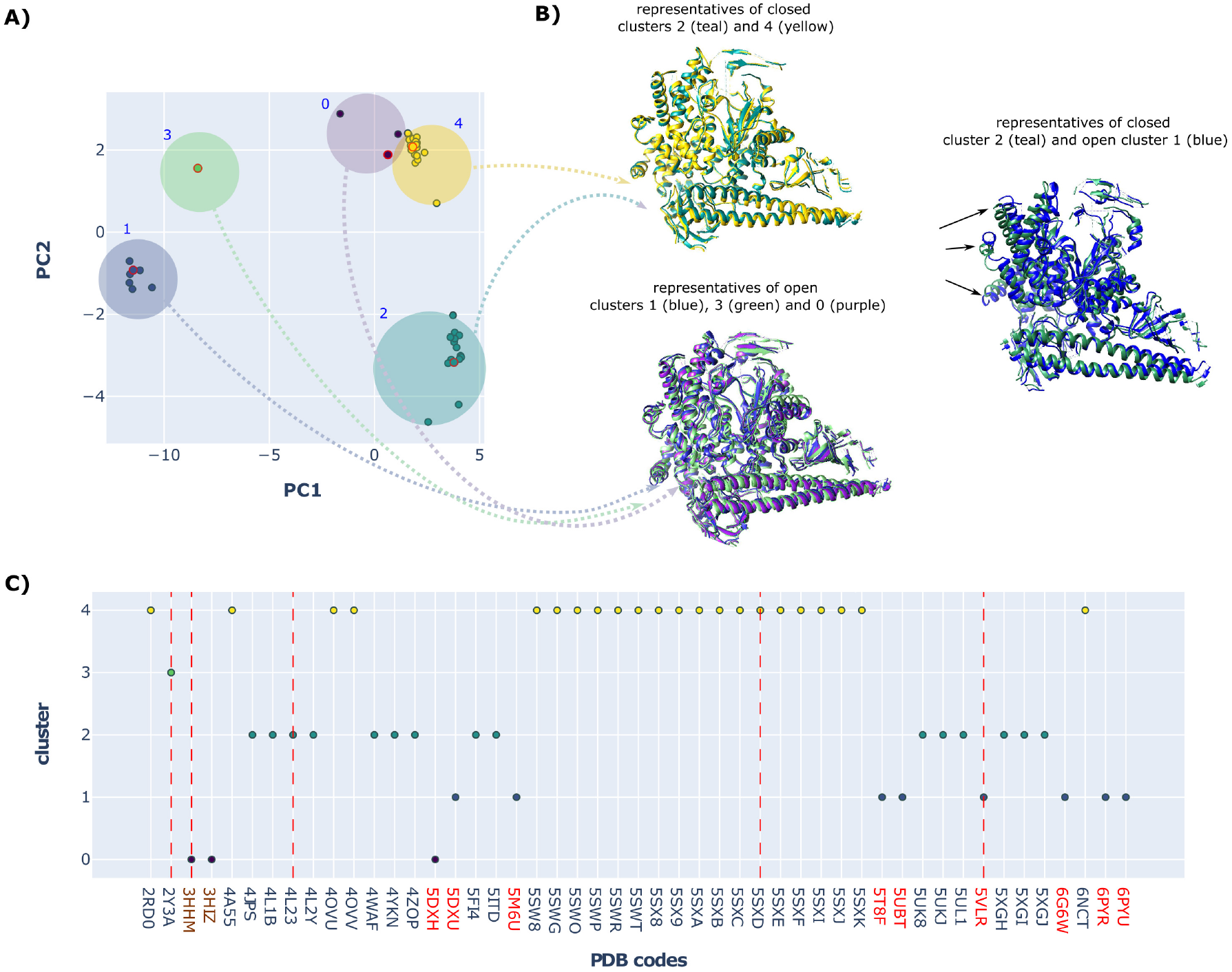
EnGens processing of the Zhang et al. dataset of PI3K crystal structures. A) Each point corresponds to a structure from Zhang et al. dataset. Points are colored based on the cluster they were assigned to, and clusters are indicated as large circles on the plot. Points extracted as cluster representatives are highlighted in red. The x and y axis represent the first and second principal components of these data. B) 3D structural models of the EnGens representatives: upper left - representatives of clusters 2 and 4 (inactive/closed states); bottom left - representatives of clusters 0, 1 and 3 (active/open states); right - comparison between the representative of the active state cluster 1 and the representative of the inactive state cluster 2 (the arrows point to the regions showing the biggest differences). C) PDB codes of the crystal structures are listed on the x axis. These codes are colored based on the conformational state identified by Zhang et al. (black - inactive/closed states; red - active/open states; brown - states active due to mutation). The EnGens cluster assignment is shown on the y axis. Red vertical lines indicate cluster representatives that were selected by EnGens (codes: 2Y3A, 3HHM, 4L23, 5SXD, 5VLR).

The dataset is clustered into five clusters. Cluster #0 contains active/open conformations of the PI3K alpha isoform (with pdb codes: 3HHM, 3HIZ and 5DXH). Cluster #1 contains eight active/open conformations of the delta isoform. Cluster #3 contains a single active/open conformation of the beta isoform (2Y3A). Clusters #2 and #4 contain the remaining 37 inactive/closed conformations of the PI3K alpha isoform. The ensemble generated by EnGens contains the following representatives: 3HHM (cluster 0), 5VLR (cluster 1), 4L23 (cluster 2), 2Y3A (cluster 3), 5SXD (cluster 4). This ensemble is structurally diverse and contains both active (3HHM, 5VLR, 2Y3A) and inactive (4L23, 5SXD) conformations. Additionally, the clusters separate the isoforms present in the dataset, namely the alpha (3HHM, 4L23, 5SXD), beta (2Y3A) and delta (5VLR) isoforms.

#### 3.1.2. PI3K-IA: MD trajectory

This experiment is based on a study by [67] involving MD simulations of a PI3K-IA (with a hotspot E545K mutation leading to its increased activity), based on multiple walkers metadynamics simulations. Galdadas et al. manually defined two collective variables: CV1 - distance between the nSH2 domain of the regulatory unit and the helical domain of the catalytic unit; CV2 - distance to a reference state where nSH2 is detached. After inspecting the free energy surface landscape as a function of CV1 and CV2, they uncovered two energy basins: one containing a conformational ensemble corresponding to an active state with the nSH2 domain detached; the other containing two distinct conformational ensembles corresponding to an alternative activation path involving nSH2 sliding around the helical domain.

We process the MD performed by Galdadas et al. with EnGens to uncover the same conformational ensembles. To featurize the trajectory we select: (1) the RMSD distance of each frame to the reference structure (first frame of the trajectory) and (2) the Cartesian coordinates of the center of mass of the helical and nSH2 domains. Next, we select SRV with a lag time of 50 to reduce the dimensionality of our input to the top 3 SRV components. We select the GMM clustering, which produces three clusters. Finally, three representative conformations are extracted. The resulting EnGens dashboard is presented in Figure 3.

**Figure 3:**
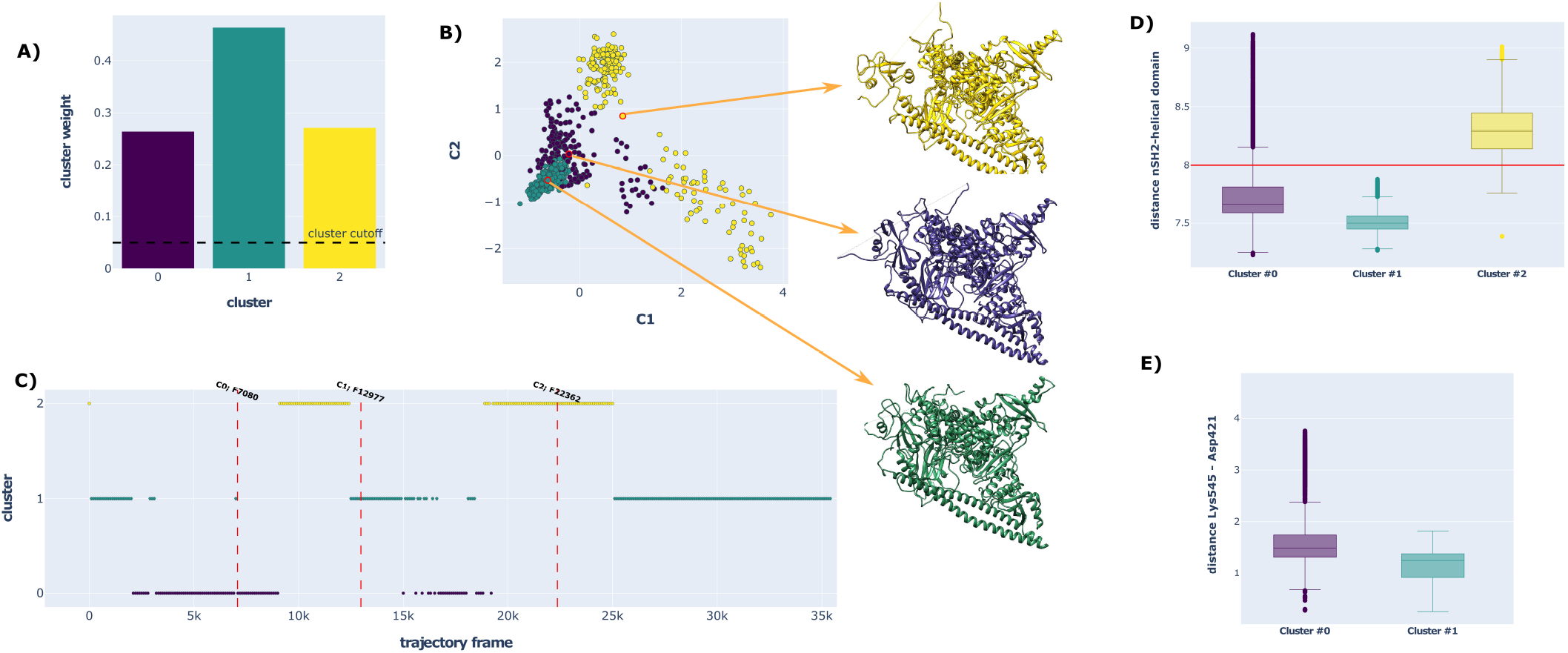
EnGens processing of the Galdadas et al. MD trajectory of PI3K. A) The proportion belonging to each cluster is plotted on the y-axis (cluster weight). Cluster indexes are listed on the x-axis. B) Two-dimensional embedding based on the components identified by the SRV method. Datapoints represent frames and are colored based on their respective cluster (same colors as in A). The 3D structural models of the three cluster representatives are shown on the right of this plot. C) The timeline view of the trajectory, where the x-axis lists the index of each frame, and the y-axis lists the corresponding cluster index. Vertical red lines highlight the representative frames extracted in the generated ensemble. D) The distance between nSH2 and the helical domain of PI3K is plotted on the y-axis. The x-axis lists the clusters. The red horizontal line represents the threshold of 8Å. E) The distance between Lys545 and Asp421 of the PI3K regulatory unit is plotted on the y-axis. The x-axis lists the clusters.

Clusters #0 and #1 contain conformations of the broad energy basin where the nSH2 domain is attached to the catalytic unit. Cluster #2 contains conformations in which PI3K is active and the nSH2 domain is detached. This is verified by plotting the minimum distance between residues of the helical domain (catalytic unit) and residues of the nSH2 domain (Figure 3D) for all cluster members. This distance stands out for members of cluster #2 and is higher than the 2 Å threshold. Clusters #0 and #1 contain the two conformational ensembles located in the same energy basin, as identified by the original paper. These clusters differ in the distance between the residues Lys545 of the helical domain and Arg421 of the nSH2 unit (Figure 3E). In conclusion, the three described clusters identified by EnGens are consistent with the three described states reported by Galdadas et al.

### 3.2. Compstatin experiment

Compstatin is a small, cyclic peptide that inhibits an immune surveillance mechanism associated with multiple diseases. Previously, we demonstrated that compstatin analogs (i.e., biochemical variants) adopt distinct conformations that ultimately affect binding affinity and inhibitor potential [68].

We applied EnGens workflows to two compstatin analogs, using two MD simulations [68]. The 4MeW and Cp10 analogs were selected because of their conformational heterogeneity. To featurize conformations, we selected backbone torsions and carbon-alpha distances. Features are then summarized using UMAP and clustered using K-means.

As a result, we retrieve several representative conformations spanning different states of these analogs (Figure 4). In particular, EnGens could accurately retrieve conformational states associated with 4WeM, namely the open *v*-shaped state, the closed *α*-shaped, and three intermediate states. These states are identified as five distinct clusters and are structurally similar to our prior observations. This demonstrates again that EnGens can reproduce results obtained with distinct methodologies. The Cp10 analog showed intriguing results. We obraine three clusters corresponding to distinct conformations, including an intermediate state. In our previous study,we could assign only two states (the opened *v*-shaped and closed *α*-shaped ones) for the Cp10 analog. To our understanding, this discrepancy is due to the limitations of our previous analysis, which only relied on RMSD calculations and visual data interpretation.

**Figure 4:**
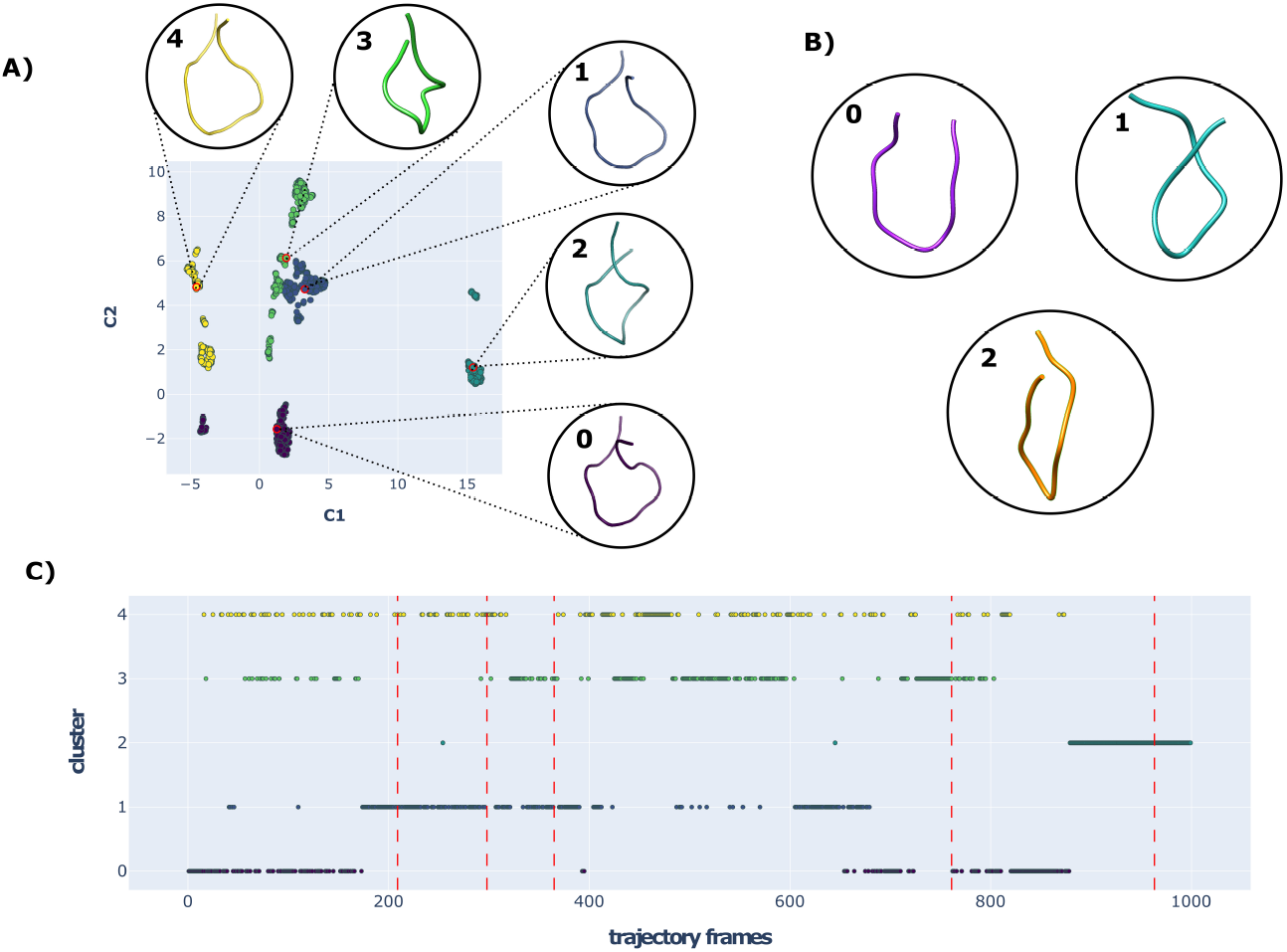
EnGens analysis of Compstatin analogs. A) UMAP visualization of the MD trajectory of the 4MeW analog. Points represent MD frames, and are colored based on their respective cluster. Cluster representatives (points highlighted in red) were selected as the closest conformation from the k-mean centroid. The 4MeW cartoon backbone for each cluster representative is presented, including a single closed state (Cluster #2), a single open state (Cluster #1), and three intermediate states (Cluster #0, #3, #4). B) The Cp10 analog: cartoon backbone visualizations of the three cluster representatives, namely the open opened, closed, and intermediate states, respectively. C) Time-oriented plot displaying the association between each MD frame (x-axis) and the clusters.

These new findings suggest that EnGens has high sensitivity and can capture rapid transitions between conformational states.

### 3.3. Nelfinavir experiment

Nelfinavir is a potent HIV-1 protease inhibitor used in adults and children. Its action mechanism involves disabling the protease from cleaving gag-pol polyprotein. However, mutations of the protease might affect the impact of Nelfinavir on patients. Using MD simulations of Nelfinavir in solution, [69] inspected its conformational space and described three minimal energy Nelfinavir conformations.

We apply EnGens to these MD trajectories of Nelfinavir. We used the Cartesian coordinates of all the atoms of Nelfinavir as the featurization. Then, we apply SRV for dimensionality reduction and GMM for clustering (Figure 5A). As a result, EnGens identifies seven clusters (Figure 5B). One cluster representative conformation coincides with the conformation described as NF-i1 and other representatives are similar to the conformations described as NF-i2 and NF-i3 in the original paper (Figure 5C), considering an RMSD under 2 Å of difference (Table 1).

**Table 1:**
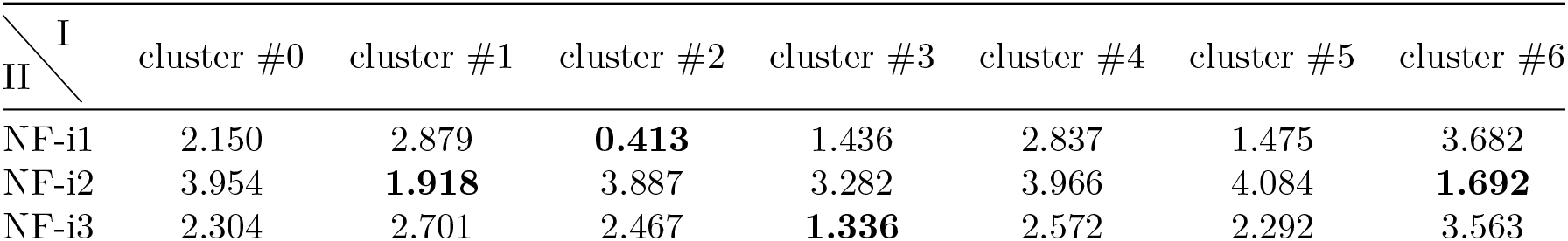
Root mean square deviation (RMSD in Å) of the EnGens cluster representatives (I) for [69] trajectory to the conformations from the original paper (II). Representatives are generated by EnGens using SRV for dimensionality reduction and GMM for clustering. RMSD values below 2Å are bolded.

**Figure 5:**
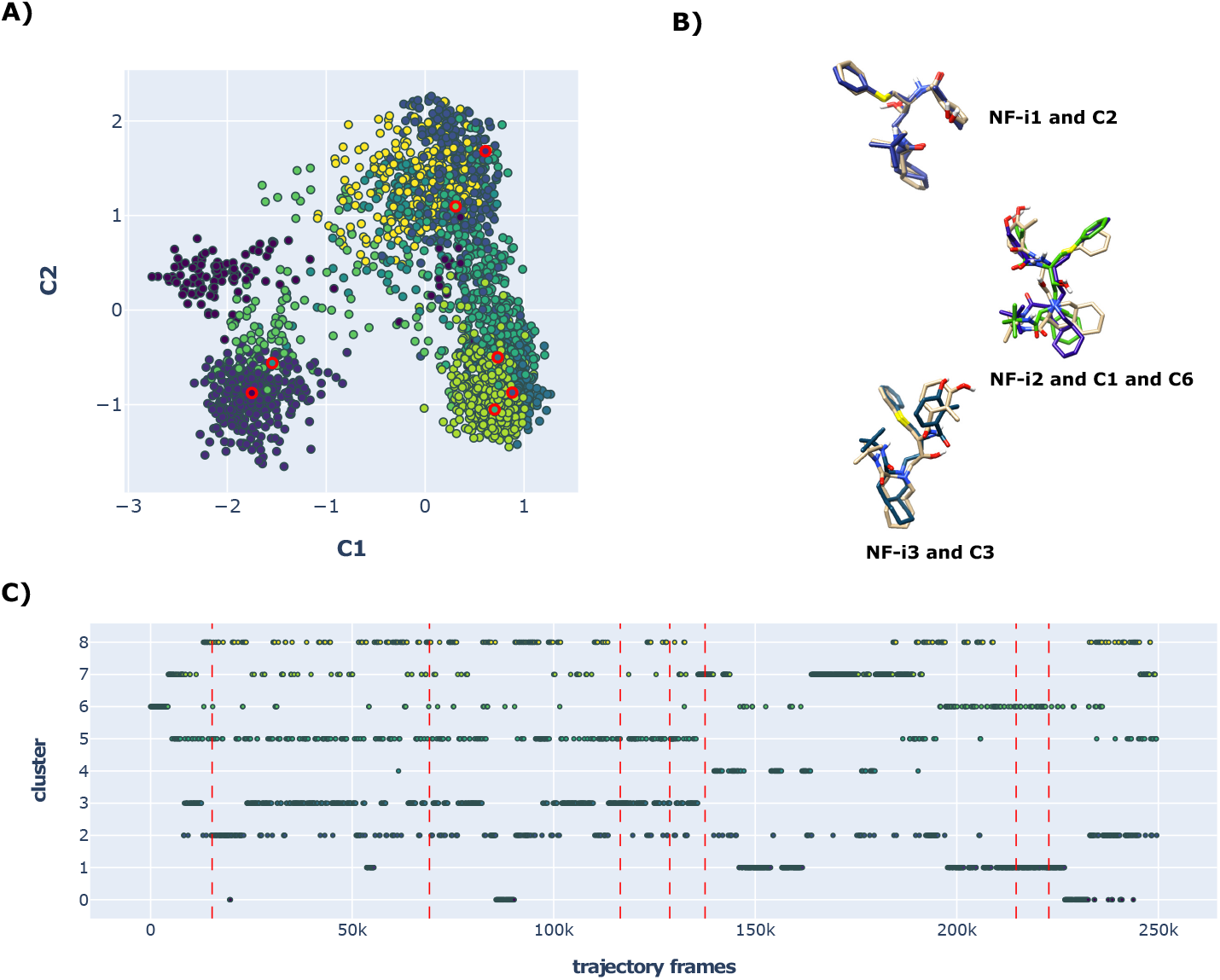
EnGens analysis of the Nelfinavir trajectory. A) Projection of all MD frames into a 2D space produced by SRV. Points are colored based on the cluster to which frames were assigned. B) Three plots showing the representative of the clusters aligned with the conformations identified by [69] (NF-i1, NF-i2, NF-i3 in pale tan color). C) Timeline of the MD trajectory showing which frame (x-axis) belongs to which cluster (y-axis).

The first conformation, NF-i1, coincides with the representative of cluster #2 (RMSD = 0.413 Å). The NF-i2 structure matches cluster #1 and #6, with RMSD of 1.918 Å and 1.692 Å respectively. However, both clusters are at the end of the trajectory (Figure 5C), where they strongly overlap, indicating that EnGens slightly refined the state corresponding to the conformation presented in the original paper. The NF-i3 conformation is to EnGens’ cluster #3, with an RMSD of 1.336 Å.

## 4. Discussion

Recent improvements in protein structure prediction tools are bringing the field of computational structural biology closer to the era of big data. One important “unsolved” task highlighted by the most recent CASP15 competition is modeling protein conformational ensembles. It is assumed that an ensemble of protein conformations will better represent the true state of a protein and will aid downstream tasks such as drug-target interaction prediction and molecular docking. Building a representative protein conformational ensemble from multiple input structures or an MD trajectory is not an easy task and many tools have been developed to tackle it. In this work, we recognized a need for a pipeline that we call EnGens.

We have evaluated the EnGens pipeline on systems of varying complexity. In each case, we recovered diverse ensembles that coincided with previously reported results. When analyzing a large protein complex such as PI3K, EnGens generated a representative ensemble containing both the active and inactive states. For the Compstatin peptide EnGens uncovered additional clusters of conformations, therefore enriching a previous study. In addition, EnGens also generated a relevant ensemble for the small drug Nelfinavir.

There are still big challenges for a pipeline of this sort. First, there are no clear guidelines on which method would perform best for a given molecular system. A number of alternative methods, each bearing its own set of hyper-parameters (Table S3), can be used to perform steps of the pipeline. We provide default values and some theoretical guidelines. For example, SRV and UMAP perform nonlinear dimensionality reduction, while TICA and PCA are linear. We thus suggest using SRV and UMAP for more complex systems where nonlinearity of features is expected. In addition, as TICA and SRV are techniques that are suitable for time-series data, they are expected to be less prone to noise resulting from fast fluctuations in the structure and should be suitable for the dynamic use-case. However, they can not be applied to the static use-case. Further theoretical analysis of some of these methods can be found in the literature [70]. Hyper-parameter optimization of the pipeline could be tackled with Bayesian optimization or other machine-learning approaches [71]. However, a wider benchmarking of these methods is necessary to evaluate the practical implications of the theory and provide good guidelines.

Second, expert knowledge of the analyzed system is still recommended for the featurization step. Some featurizations are generic, such as the pairwise residue distances that we applied to Compstatin. Others, such as the distance between the nSH2 domain and the helical domain of PI3K stem from a good understanding of the underlying system. Efforts have been made to automate this step. For the dynamic use-case new breakthroughs such as VAC (Variational Approach to Conformational dynamics) [72, 73] and VAMP (Variational Approach for Learning Markov Processes) [72] provide metrics to quantify the quality of featurization. Such metrics can be optimized using machine learning approaches to determine the most suitable featurization. However, these methods are highly dependent on the quality of the provided input MD data and are sensitive to different hyperparameters. Engineering features manually is still a widely used practice.

Third, we lack large standardized benchmarks and metrics for generating conformational ensembles. To avoid the hurdles we faced in this work, the community would greatly benefit from a public database collecting both static and dynamic datasets of protein conformations for which the representative conformational ensembles are known. Another problem is the lack of standardized metrics to compare the uncovered conformational states. Although RMSD is widely used to compare protein conformations, there are currently no equivalent standardized metrics for comparing two conformational ensembles. That is why our evaluation of EnGens is mostly qualitative and descriptive (e.g. determining if EnGens uncovered the active and inactive conformational states of PI3K).

These challenges will become more pressing as the field moves towards big data analysis to study protein flexibility. EnGens provides easy access to existing algorithms and can serve as a platform for the rapid development of new algorithms addressing these challenges.

## 5. Conclusion

EnGens is a novel tool for the end-to-end processing of large protein structural datasets with the aim of generating and analyzing representative protein conformational ensembles. EnGens unifies widely used Python libraries (PyEmma, deeptime, mdtraj, UMAP, sklearn, plotly, etc.) under one Docker image and provides interactive visualizations along with extensive examples of the pipeline in Jupyter Notebook workflows. For advanced users, we provide a Python package. Our code is open source and accessible through a github repository (https://github.com/KavrakiLab/EnGens). We showcased how EnGens can be used to automate ensemble generation using examples from the literature. EnGens ensembles can be useful for many downstream tasks related to drug discovery such as molecular docking and drug-target interaction prediction. Additionally, EnGens can serve as a platform for further algorithmic development. Overall, we see the EnGens pipeline becoming part of many new efforts to utilize the structural data generated by novel structure prediction tools.

## 6. Author contributions statement

A.C., M.M.R., D.D., A.F.F., M.V.F., C.C., G.Z., and D.A.A. conceived the experiments. A.C., A.F.F., H.K., and M.V.F. conducted the experiments. A.C., A.F.F., and M.V.F. wrote the manuscript, A.C., M.M.R., D.D., A.F.F., M.V.F., C.C., G.Z., D.A.A. and L.K. analyzed the results. All authors reviewed the manuscript.

## 7. Acknowledgments

The authors would like to thank colleagues from Kavraki Lab for many helpful discussions.

## References

[1] A. Kessel, N. Ben-Tal, : Structure, Function, and Motion, Second Edition, 2nd Edition, Chapman and Hall/CRC, New York, 2018. doi:10.1201/9781315113876.

[2] R. Nussinov, C.-J. Tsai, Allostery in Disease and in Drug Discovery, Cell 153 (2) (2013) 293–305. doi:10.1016/j.cell.2013.03.034. URL https://www.sciencedirect.com/science/article/pii/S0092867413003917

[3] A. E. Todd, C. A. Orengo, J. M. Thornton, Plasticity of enzyme active sites, Trends in Biochemical Sciences 27 (8) (2002) 419–426, publisher: Elsevier. doi:10.1016/S0968-0004(02)02158-8. URL https://www.cell.com/trends/biochemical-sciences/abstract/S0968-0004(02)02158-8

[4] C.-L. Tsou, Active Site Flexibility in Enzyme Catalysisa, Annals of the New York Academy of Sciences 864 (1) (1998) 1–8, _eprint: https://onlinelibrary.wiley.com/doi/pdf/10.1111/j.1749-6632.1998.tb10282.x. doi:10.1111/j.1749-6632.1998.tb10282.x. URL https://onlinelibrary.wiley.com/doi/abs/10.1111/j.1749-6632.1998.tb10282.

[5] Y.-Z. Weng, D. T.-H. Chang, Y.-F. Huang, C.-W. Lin, A study on the flexibility of enzyme active sites, BMC Bioinformatics 12 (1) (2011) S32. doi:10.1186/1471-2105-12-S1-S32. URL https://doi.org/10.1186/1471-2105-12-S1-S32

[6] A. F. Dishman, B. F. Volkman, Unfolding the Mysteries of Protein Metamorphosis, ACS Chemical Biology 13 (6) (2018) 1438–1446. doi:10.1021/acschembio.8b00276. URL https://www.ncbi.nlm.nih.gov/pmc/articles/PMC6007232/

[7] A. F. Dishman, B. F. Volkman, Design and discovery of metamorphic proteins, Current Opinion in Structural Biology 74 (2022) 102380. doi:10.1016/j.sbi.2022.102380. URL https://www.sciencedirect.com/science/article/pii/S0959440X22000598

[8] M. Lella, R. Mahalakshmi, Metamorphic Proteins: Emergence of Dual Protein Folds from One Primary Sequence, Biochemistry 56 (24) (2017) 2971–2984, publisher: American Chemical Society. doi:10.1021/acs.biochem.7b00375. URL https://doi.org/10.1021/acs.biochem.7b00375

[9] A. K. Kim, L. L. Porter, Functional and Regulatory Roles of Fold-Switching Proteins, Structure 29 (1) (2021) 6–14. doi:10.1016/j.str.2020.10.006. URL https://www.sciencedirect.com/science/article/pii/S0969212620303786

[10] V. N. Uversky, p53 Proteoforms and Intrinsic Disorder: An Illustration of the Protein Structure–Function Continuum Concept, International Journal of Molecular Sciences 17 (11) (2016) 1874. doi:10.3390/ijms17111874. URL https://www.ncbi.nlm.nih.gov/pmc/articles/PMC5133874/

[11] K. Henzler-Wildman, D. Kern, Dynamic personalities of proteins, Nature 450 (7172) (2007) 964–972, number: 7172 Publisher: Nature Publishing Group. doi:10.1038/nature06522. URL https://www.nature.com/articles/nature06522

[12] H. Frauenfelder, S. G. Sligar, P. G. Wolynes, The Energy Landscapes and Motions of Proteins, Science 254 (5038) (1991) 1598–1603, publisher: American Association for the Advancement of Science. doi:10.1126/science.1749933. URL https://www.science.org/doi/10.1126/science.1749933

[13] S. Kumar, B. Ma, C.-J. Tsai, N. Sinha, R. Nussinov, Folding and binding cascades: Dynamic landscapes and population shifts, Protein Science 9 (1) (2000) 10–19, publisher: Cambridge University Press. doi:10.1110/ps.9.1.10. URL https://www.cambridge.org/core/journals/protein-science/article/abs/folding-and-binding-cascades-dynamic-landscapes-and-population-shifts/C7CFF2407012931BA6C0D1A898C9D922

[14] J. N. Onuchic, Z. Luthey-Schulten, P. G. Wolynes, Theory of protein folding: the energy landscape perspective, Annual Review of Physical Chemistry 48 (1997) 545–600. doi:10.1146/annurev.physchem.48.1.545.

[15] J. Jumper, R. Evans, A. Pritzel, T. Green, M. Figurnov, O. Ronneberger, K. Tunyasuvunakool, R. Bates, A. Žídek, A. Potapenko, A. Bridgland, C. Meyer, S. A. A. Kohl, A. J. Ballard, A. Cowie, B. Romera-Paredes, S. Nikolov, R. Jain, J. Adler, T. Back, S. Petersen, D. Reiman, E. Clancy, M. Zielinski, M. Steinegger, M. Pacholska, T. Berghammer, S. Bodenstein, D. Silver, O. Vinyals, A. W. Senior, K. Kavukcuoglu, P. Kohli, D. Hassabis, Highly accurate protein structure prediction with AlphaFold, Nature 596 (7873) (2021) 583–589, number: 7873 Publisher: Nature Publishing Group. doi:10.1038/s41586-021-03819-2. URL https://www.nature.com/articles/s41586-021-03819-2

[16] M. Baek, F. DiMaio, I. Anishchenko, J. Dauparas, S. Ovchinnikov, G. R. Lee, J. Wang, Q. Cong, L. N. Kinch, R. D. Schaeffer, C. Millán, H. Park, C. Adams, C. R. Glassman, A. DeGiovanni, J. H. Pereira, A. V. Rodrigues, A. A. van Dijk, A. C. Ebrecht, D. J. Opperman, T. Sagmeister, C. Buhlheller, T. Pavkov-Keller, M. K. Rathinaswamy, U. Dalwadi, C. K. Yip, J. E. Burke, K. C. Garcia, N. V. Grishin, P. D. Adams, R. J. Read, D. Baker, Accurate prediction of protein structures and interactions using a three-track neural network, Science 373 (6557) (2021) 871–876, publisher: American Association for the Advancement of Science. doi:10.1126/science.abj8754. URL https://www.science.org/doi/10.1126/science.abj8754

[17] Z. Lin, H. Akin, R. Rao, B. Hie, Z. Zhu, W. Lu, N. Smetanin, R. Verkuil, O. Kabeli, Y. Shmueli, A. d. S. Costa, M. Fazel-Zarandi, T. Sercu, S. Candido, A. Rives, Evolutionary-scale prediction of atomic level protein structure with a language model, pages: 2022.07.20.500902 Section: New Results (Dec. 2022). doi:10.1101/2022.07.20.500902. URL https://www.biorxiv.org/content/10.1101/2022.07.20.500902v3

[18] M. S. Barhaghi, B. Crawford, G. Schwing, D. J. Hardy, J. E. Stone, L. Schwiebert, J. Potoff, E. Tajkhorshid, py-MCMD: Python Software for Performing Hybrid Monte Carlo/Molecular Dynamics Simulations with GOMC and NAMD, Journal of Chemical Theory and Computation 18 (8) (2022) 4983–4994, publisher: American Chemical Society. doi:10.1021/acs.jctc.1c00911. URL https://doi.org/10.1021/acs.jctc.1c00911

[19] M. J. Abraham, T. Murtola, R. Schulz, S. Páll, J. C. Smith, B. Hess, E. Lindahl, GROMACS: High performance molecular simulations through multi-level parallelism from laptops to supercomputers, SoftwareX 1-2 (2015) 19–25. doi:10.1016/j.softx.2015.06.001. URL https://www.sciencedirect.com/science/article/pii/S2352711015000059

[20] P. Eastman, M. S. Friedrichs, J. D. Chodera, R. J. Radmer, C. M. Bruns, J. P. Ku, K. A. Beauchamp, T. J. Lane, L.-P. Wang, D. Shukla, T. Tye, M. Houston, T. Stich, C. Klein, M. R. Shirts, V. S. Pande, OpenMM 4: A Reusable, Extensible, Hardware Independent Library for High Performance Molecular Simulation, Journal of Chemical Theory and Computation 9 (1) (2013) 461–469, publisher: American Chemical Society. doi:10.1021/ct300857j. URL https://doi.org/10.1021/ct300857j

[21] B. E. Husic, N. E. Charron, D. Lemm, J. Wang, A. Pérez, M. Majewski, A. Krämer, Y. Chen, S. Olsson, G. de Fabritiis, F. Noé, C. Clementi, Coarse graining molecular dynamics with graph neural networks, The Journal of Chemical Physics 153 (19) (2020) 194101, publisher: American Institute of Physics. doi:10.1063/5.0026133. URL https://aip.scitation.org/doi/full/10.1063/5.0026133

[22] J. Hénin, T. Lelièvre, M. R. Shirts, O. Valsson, L. Delemotte, Enhanced sampling methods for molecular dynamics simulations, Living Journal of Computational Molecular Science 4 (1), arXiv:2202.04164 [cond-mat, physics:physics]. doi:10.33011/livecoms.4.1.1583. URL http://arxiv.org/abs/2202.04164

[23] J.-h. Peng, W. Wang, Y.-q. Yu, H.-l. Gu, X. Huang, Clustering algorithms to analyze molecular dynamics simulation trajectories for complex chemical and biological systems, Chinese Journal of Chemical Physics 31 (4) (2018) 404–420, publisher: American Institute of Physics. doi: 10.1063/1674-0068/31/cjcp1806147. URL https://cps.scitation.org/doi/10.1063/1674-0068/31/cjcp1806147

[24] S. Hall-Swan, D. Devaurs, M. M. Rigo, D. A. Antunes, L. E. Kavraki, G. Zanatta, DINC-COVID: A webserver for ensemble docking with flexible SARS-CoV-2 proteins, Computers in Biology and Medicine 139 (2021) 104943. doi:10.1016/j.compbiomed.2021.104943. URL https://www.ncbi.nlm.nih.gov/pmc/articles/PMC8518241/

[25] A. Kannan, A. N. Naganathan, Ensemble origins and distance-dependence of long-range mutational effects in proteins, iScience 25 (10) (2022) 105181. doi:10.1016/j.isci.2022.105181. URL https://www.sciencedirect.com/science/article/pii/S2589004222014535

[26] J. R. Abella, D. Antunes, K. Jackson, G. Lizée, C. Clementi, L. E. Kavraki, Markov state modeling reveals alternative unbinding pathways for peptide–MHC complexes, Proceedings of the National Academy of Sciences 117 (48) (2020) 30610–30618, publisher: Proceedings of the National Academy of Sciences. doi:10.1073/pnas.2007246117. URL https://www.pnas.org/doi/10.1073/pnas.2007246117

[27] M. C. Chan, D. Shukla, Markov state modeling of membrane transport proteins, Journal of Structural Biology 213 (4) (2021) 107800. doi:10.1016/j.jsb.2021.107800.

[28] wwPDB consortium, Protein Data Bank: the single global archive for 3D macromolecular structure data, Nucleic Acids Research 47 (D1) (2019) D520–D528. doi:10.1093/nar/gky949. URL https://doi.org/10.1093/nar/gky949

[29] S. K. Burley, C. Bhikadiya, C. Bi, S. Bittrich, L. Chen, G. V. Crichlow, C. H. Christie, K. Dalenberg, L. Di Costanzo, J. M. Duarte, S. Dutta, Z. Feng, S. Ganesan, D. S. Goodsell, S. Ghosh, R. K. Green, V. Guranović, D. Guzenko, B. P. Hudson, C. Lawson, Y. Liang, R. Lowe, H. Namkoong, E. Peisach, I. Persikova, C. Randle, A. Rose, Y. Rose, A. Sali, J. Segura, M. Sekharan, C. Shao, Y.-P. Tao, M. Voigt, J. Westbrook, J. Y. Young, C. Zardecki, M. Zhuravleva, RCSB Protein Data Bank: powerful new tools for exploring 3D structures of biological macromolecules for basic and applied research and education in fundamental biology, biomedicine, biotechnology, bioengineering and energy sciences, Nucleic Acids Research 49 (D1) (2021) D437–D451. doi:10.1093/nar/gkaa1038. URL https://doi.org/10.1093/nar/gkaa1038

[30] M. Varadi, S. Anyango, M. Deshpande, S. Nair, C. Natassia, G. Yordanova, D. Yuan, O. Stroe, G. Wood, A. Laydon, A. Žídek, T. Green, K. Tunyasuvunakool, S. Petersen, J. Jumper, E. Clancy, R. Green, A. Vora, M. Lutfi, M. Figurnov, A. Cowie, N. Hobbs, P. Kohli, G. Kleywegt, E. Birney, D. Hassabis, S. Velankar, AlphaFold Protein Structure Database: massively expanding the structural coverage of protein-sequence space with high-accuracy models, Nucleic Acids Research 50 (D1) (2022) D439–D444. doi:10.1093/nar/gkab1061. URL https://doi.org/10.1093/nar/gkab1061

[31] Y. Takei, T. Ishida, How to select the best model from AlphaFold2 structures?, preprint, Bioinformatics (Apr. 2022). doi:10.1101/2022.04.05.487218. URL http://biorxiv.org/lookup/doi/10.1101/2022.04.05.487218

[32] A. Warshel, M. Levitt, Theoretical studies of enzymic reactions: Dielectric, electrostatic and steric stabilization of the carbonium ion in the reaction of lysozyme, Journal of Molecular Biology 103 (2) (1976) 227–249. doi:10.1016/0022-2836(76)90311-9. URL https://www.sciencedirect.com/science/article/pii/0022283676903119

[33] J. C. Phillips, D. J. Hardy, J. D. C. Maia, J. E. Stone, J. V. Ribeiro, R. C. Bernardi, R. Buch, G. Fiorin, J. Hénin, W. Jiang, R. McGreevy, M. C. R. Melo, B. K. Radak, R. D. Skeel, A. Singharoy, Y. Wang, B. Roux, A. Aksimentiev, Z. Luthey-Schulten, L. V. Kalé, K. Schulten, C. Chipot, E. Tajkhorshid, Scalable molecular dynamics on CPU and GPU architectures with NAMD, The Journal of Chemical Physics 153 (4) (2020) 044130, publisher: American Institute of Physics. doi:10.1063/5.0014475. URL https://aip.scitation.org/doi/abs/10.1063/5.0014475

[34] H. Bekker, H. Berendsen, E. Dijkstra, S. Achterop, R. Vondrumen, D. Van- Derspoel, A. Sijbers, H. Keegstra, M. Renardus, Gromacs - A PARALLEL COMPUTER FOR MOLECULAR-DYNAMICS SIMULATIONS: 4th International Conference on Computational Physics (PC 92), PHYSICS COMPUTING ’92 (1993) 252–256Place: SINGAPORE Publisher: World Scientific Publishing.

[35] H. J. C. Berendsen, D. van der Spoel, R. van Drunen, GROMACS: A message-passing parallel molecular dynamics implementation, Computer Physics Communications 91 (1) (1995) 43–56. doi:10.1016/0010-4655(95)00042-E. URL https://www.sciencedirect.com/science/article/pii/001046559500042E

[36] R. Salomon-Ferrer, D. A. Case, R. C. Walker, An overview of the Amber biomolecular simulation package, WIREs Computational Molecular Science 3 (2) (2013) 198–210, _eprint: https://onlinelibrary.wiley.com/doi/pdf/10.1002/wcms.1121. doi:10.1002/wcms.1121. URL https://onlinelibrary.wiley.com/doi/abs/10.1002/wcms.1121

[37] B. Brooks, C. Brooks, A. MacKerell, L. Nilsson, R. Petrella, B. Roux, Y. Won, G. Archontis, C. Bartels, S. Boresch, A. Caflisch, L. Caves, Q. Cui, A. Dinner, M. Feig, S. Fischer, J. Gao, M. Hodoscek, W. Im, K. Kuczera, T. Lazaridis, J. Ma, V. Ovchinnikov, E. Paci, R. Pastor, C. Post, J. Pu, M. Schaefer, B. Tidor, R. M. Venable, H. L. Woodcock, X. Wu, W. Yang, D. York, M. Karplus, CHARMM: The Biomolecular Simulation Program, Journal of computational chemistry 30 (10) (2009) 1545–1614. doi:10.1002/jcc.21287. URL https://www.ncbi.nlm.nih.gov/pmc/articles/PMC2810661/

[38] P. Eastman, J. Swails, J. D. Chodera, R. T. McGibbon, Y. Zhao, K. A. Beauchamp, L.-P. Wang, A. C. Simmonett, M. P. Harrigan, C. D. Stern, R. P. Wiewiora, B. R. Brooks, V. S. Pande, OpenMM 7: Rapid development of high performance algorithms for molecular dynamics, PLoS computational biology 13 (7) (2017) e1005659. doi:10.1371/journal.pcbi.1005659.

[39] J.-H. Prinz, H. Wu, M. Sarich, B. Keller, M. Senne, M. Held, J. D. Chodera, C. Schütte, F. Noé, Markov models of molecular kinetics: Generation and validation, The Journal of Chemical Physics 134 (17) (2011) 174105, publisher: American Institute of Physics. doi: 10.1063/1.3565032. URL https://aip.scitation.org/doi/10.1063/1.3565032

[40] P. J. A. Cock, T. Antao, J. T. Chang, B. A. Chapman, C. J. Cox, A. Dalke, I. Friedberg, T. Hamelryck, F. Kauff, B. Wilczynski, M. J. L. de Hoon, Biopython: freely available Python tools for computational molecular biology and bioinformatics, Bioinformatics 25 (11) (2009) 1422–1423. doi:10.1093/bioinformatics/btp163. URL https://doi.org/10.1093/bioinformatics/btp163

[41] M. K. Scherer, B. Trendelkamp-Schroer, F. Paul, G. Pérez-Hernández, M. Hoffmann, N. Plattner, C. Wehmeyer, J.-H. Prinz, F. Noé, PyEMMA 2: A Software Package for Estimation, Validation, and Analysis of Markov Models, Journal of Chemical Theory and Computation 11 (11) (2015) 5525–5542, publisher: American Chemical Society. doi:10.1021/acs.jctc.5b00743. URL https://doi.org/10.1021/acs.jctc.5b00743

[42] R. McGibbon, K. Beauchamp, M. Harrigan, C. Klein, J. Swails, C. Hernández, C. Schwantes, L.-P. Wang, T. Lane, V. Pande, MDTraj: A Modern Open Library for the Analysis of Molecular Dynamics Trajectories, Biophysical Journal 109 (8) (2015) 1528–1532. doi:10.1016/j.bpj.2015.08.015. URL https://www.ncbi.nlm.nih.gov/pmc/articles/PMC4623899/

[43] M. Hoffmann, M. Scherer, T. Hempel, A. Mardt, B. d. Silva, B. E. Husic, S. Klus, H. Wu, N. Kutz, S. L. Brunton, F. Noé, Deeptime: a Python library for machine learning dynamical models from time series data, Machine Learning: Science and Technology 3 (1) (2021) 015009, publisher: IOP Publishing. doi:10.1088/2632-2153/ac3de0. URL https://dx.doi.org/10.1088/2632-2153/ac3de0

[44] F. Pedregosa, G. Varoquaux, A. Gramfort, V. Michel, B. Thirion, O. Grisel, M. Blondel, P. Prettenhofer, R. Weiss, V. Dubourg, J. Vanderplas, A. Passos, D. Cournapeau, M. Brucher, M. Perrot, Duchesnay, Scikit-learn: Machine Learning in Python, Journal of Machine Learning Research 12 (85) (2011) 2825–2830. URL http://jmlr.org/papers/v12/pedregosa11a.html

[45] F. Trozzi, X. Wang, P. Tao, UMAP as a Dimensionality Reduction Tool for Molecular Dynamics Simulations of Biomacromolecules: A Comparison Study, The Journal of Physical Chemistry B 125 (19) (2021) 5022–5034, publisher: American Chemical Society. doi:10.1021/acs.jpcb.1c02081. URL https://doi.org/10.1021/acs.jpcb.1c02081

[46] W. Chen, H. Sidky, A. L. Ferguson, Nonlinear discovery of slow molecular modes using state-free reversible VAMPnets, The Journal of Chemical Physics 150 (21) (2019) 214114, publisher: American Institute of Physics. doi:10.1063/1.5092521. URL https://aip.scitation.org/doi/10.1063/1.5092521

[47] A. F. Ángyán, B. Szappanos, A. Perczel, Z. Gáspári, CoNSEnsX: an ensemble view of protein structures and NMR-derived experimental data, BMC Structural Biology 10 (1) (2010) 39. doi:10.1186/1472-6807-10-39. URL https://doi.org/10.1186/1472-6807-10-39

[48] M. Vögele, N. J. Thomson, S. T. Truong, J. McAvity, U. Zachariae, R. O. Dror, Systematic Analysis of Biomolecular Conformational Ensembles with PENSA, arXiv:2212.02714 [physics, q-bio] (Dec. 2022). doi:10.48550/arXiv.2212.02714. URL http://arxiv.org/abs/2212.02714

[49] M. Vögele, N. J. Thomson, J. McAvity, Mick, drorlab/pensa: PENSA 0.2.8 (Feb. 2022). doi:10.5281/zenodo.6256993. URL https://zenodo.org/record/6256993

[50] A. Bakan, L. M. Meireles, I. Bahar, ProDy: Protein Dynamics Inferred from Theory and Experiments, Bioinformatics 27 (11) (2011) 1575–1577. doi:10.1093/bioinformatics/btr168. URL https://doi.org/10.1093/bioinformatics/btr168

[51] S. Zhang, J. M. Krieger, Y. Zhang, C. Kaya, B. Kaynak, K. Mikulska-Ruminska, P. Doruker, H. Li, I. Bahar, ProDy 2.0: increased scale and scope after 10 years of protein dynamics modelling with Python, Bioinformatics 37 (20) (2021) 3657–3659. doi:10.1093/bioinformatics/btab187. URL https://doi.org/10.1093/bioinformatics/btab187

[52] K. Pearson, LIII. On lines and planes of closest fit to systems of points in space, The London, Edinburgh, and Dublin Philosophical Magazine and Journal of Science 2 (11) (1901) 559–572, publisher: Taylor & Francis _eprint: https://doi.org/10.1080/14786440109462720. doi: 10.1080/14786440109462720. URL https://doi.org/10.1080/14786440109462720

[53] G. Pérez-Hernández, F. Paul, T. Giorgino, G. De Fabritiis, F. Noé, Identification of slow molecular order parameters for Markov model construction, The Journal of Chemical Physics 139 (1) (2013) 015102, publisher: American Institute of Physics. doi:10.1063/1.4811489. URL https://aip.scitation.org/doi/10.1063/1.4811489

[54] C. R. Schwantes, V. S. Pande, Modeling Molecular Kinetics with tICA and the Kernel Trick, Journal of Chemical Theory and Computation 11 (2) (2015) 600–608, publisher: American Chemical Society. doi:10.1021/ct5007357. URL https://doi.org/10.1021/ct5007357

[55] B. E. Husic, V. S. Pande, Markov State Models: From an Art to a Science, Journal of the American Chemical Society 140 (7) (2018) 2386–2396, publisher: American Chemical Society. doi:10.1021/jacs.7b12191. URL https://doi.org/10.1021/jacs.7b12191

[56] M. Bernetti, M. Masetti, M. Recanatini, R. E. Amaro, A. Cavalli, An Integrated Markov State Model and Path Metadynamics Approach To Characterize Drug Binding Processes, Journal of Chemical Theory and Computation 15 (10) (2019) 5689–5702, publisher: American Chemical Society. doi:10.1021/acs.jctc.9b00450. URL https://doi.org/10.1021/acs.jctc.9b00450

[57] F. Murtagh, P. Contreras, Algorithms for hierarchical clustering: an overview, WIREs Data Mining and Knowledge Discovery 2 (1) (2012) 86–97, _eprint: https://onlinelibrary.wiley.com/doi/pdf/10.1002/widm.53. doi:10.1002/widm.53. URL https://onlinelibrary.wiley.com/doi/abs/10.1002/widm.53

[58] J. A. Hartigan, M. A. Wong, Algorithm AS 136: A K-Means Clustering Algorithm, Journal of the Royal Statistical Society. Series C (Applied Statistics) 28 (1) (1979) 100–108, publisher: [Wiley, Royal Statistical Society]. doi:10.2307/2346830. URL https://www.jstor.org/stable/2346830

[59] B. Lindsay, G. L. McLachlan, K. E. Basford, M. Dekker, Mixture Models: Inference and Applications to Clustering., in: Journal of the American Statistical Association, Vol. 84, 1989, p. 337, iSSN: 01621459 Issue: 405 Journal Abbreviation: Journal of the American Statistical Association. doi:10.2307/2289892. URL https://www.jstor.org/stable/2289892?origin=crossref

[60] M. Martini, M. C. De Santis, L. Braccini, F. Gulluni, E. Hirsch, PI3K/AKT signaling pathway and cancer: an updated review, Annals of Medicine 46 (6) (2014) 372–383, publisher: Taylor & Francis _eprint: https://doi.org/10.3109/07853890.2014.912836. doi:10.3109/07853890.2014.912836. URL https://doi.org/10.3109/07853890.2014.912836

[61] J. Yu, C. Wjasow, J. M. Backer, Regulation of the p85/p110 Phosphatidylinositol 3-Kinase: DISTINCT ROLES FOR THE N-TERMINAL AND C-TERMINAL SH2 DOMAINS*, Journal of Biological Chemistry 273 (46) (1998) 30199–30203. doi:10.1074/jbc.273.46.30199. URL https://www.sciencedirect.com/science/article/pii/S0021925819592225

[62] M. S. Miller, O. Schmidt-Kittler, D. M. Bolduc, E. T. Brower, D. Chaves-Moreira, M. Allaire,K. W. Kinzler, I. G. Jennings, P. E. Thompson, P. A. Cole, L. M. Amzel, B. Vogelstein, S. B. Gabelli, Structural basis of nSH2 regulation and lipid binding in PI3K, Oncotarget 5 (14) (2014) 5198–5208. doi:10.18632/oncotarget.2263. URL https://www.oncotarget.com/lookup/doi/10.18632/oncotarget.2263

[63] T. C. Buckles, B. P. Ziemba, G. R. Masson, R. L. Williams, J. J. Falke, Single-Molecule Study Reveals How Receptor and Ras Synergistically Activate PI3K and PIP3 Signaling, Biophysical Journal 113 (11) (2017) 2396–2405. doi:10.1016/j.bpj.2017.09.018. URL https://www.sciencedirect.com/science/article/pii/S0006349517310305

[64] R. T. Nolte, M. J. Eck, J. Schlessinger, S. E. Shoelson, S. C. Harrison, Crystal structure of the PI 3-kinase p85 amino-terminal SH2 domain and its phosphopeptide complexes, Nature Structural Biology 3 (4) (1996) 364–374, number: 4 Publisher: Nature Publishing Group. doi:10.1038/nsb0496-364. URL https://www.nature.com/articles/nsb0496-364

[65] O. Vadas, J. E. Burke, X. Zhang, A. Berndt, R. L. Williams, Structural Basis for Activation and Inhibition of Class I Phosphoinositide 3-Kinases, Science Signaling 4 (195) (2011) re2–re2, publisher: American Association for the Advancement of Science. doi:10.1126/scisignal.2002165. URL https://www.science.org/doi/full/10.1126/scisignal.2002165

[66] M. Zhang, H. Jang, R. Nussinov, Structural Features that Distinguish Inactive and Active PI3K Lipid Kinases, Journal of Molecular Biology 432 (22) (2020) 5849–5859. doi:10.1016/j.jmb.2020.09.002. URL https://www.sciencedirect.com/science/article/pii/S0022283620305325

[67] I. Galdadas, F. L. Gervasio, Z. Cournia, Unravelling the effect of the E545K mutation on PI3K kinase, Chemical Science 11 (13) (2020) 3511–3515, publisher: The Royal Society of Chemistry. doi:10.1039/C9SC05903B. URL https://pubs.rsc.org/en/content/articlelanding/2020/sc/c9sc05903b

[68] D. Devaurs, D. A. Antunes, L. E. Kavraki, Computational analysis of complement inhibitor compstatin using molecular dynamics, Journal of Molecular Modeling 26 (9) (2020) 231. doi:10.1007/s00894-020-04472-8. URL https://doi.org/10.1007/s00894-020-04472-8

[69] D. A. Antunes, M. M. Rigo, M. Sinigaglia, R. M. d. Medeiros, D. M. Junqueira, S. E. M. Almeida, G. F. Vieira, New Insights into the In Silico Prediction of HIV Protease Resistance to Nelfinavir, PLOS ONE 9 (1) (2014) e87520, publisher: Public Library of Science. doi: 10.1371/journal.pone.0087520. URL https://journals.plos.org/plosone/article?id=10.1371/journal.pone.0087520

[70] A. Glielmo, B. E. Husic, A. Rodriguez, C. Clementi, F. Noé, A. Laio, Unsupervised Learning Methods for Molecular Simulation Data, Chemical Reviews 121 (16) (2021) 9722–9758. doi: 10.1021/acs.chemrev.0c01195. URL https://pubs.acs.org/doi/10.1021/acs.chemrev.0c01195

[71] Y. Lee, C. Chamzas, L. E. Kavraki, Adaptive Experience Sampling for Motion Planning Using the Generator-Critic Framework, IEEE Robotics and Automation Letters 7 (4) (2022) 9437–9444, conference Name: IEEE Robotics and Automation Letters. doi:10.1109/LRA.2022.3191803.

[72] H. Wu, F. Noé, Variational Approach for Learning Markov Processes from Time Series Data, Journal of Nonlinear Science 30 (1) (2020) 23–66. doi:10.1007/s00332-019-09567-y. URL https://doi.org/10.1007/s00332-019-09567-y

[73] C. Lorpaiboon, E. H. Thiede, R. J. Webber, J. Weare, A. R. Dinner, Integrated Variational Approach to Conformational Dynamics: A Robust Strategy for Identifying Eigenfunctions of Dynamical Operators, The Journal of Physical Chemistry B 124 (42) (2020) 9354–9364, publisher: American Chemical Society. doi:10.1021/acs.jpcb.0c06477. URL https://doi.org/10.1021/acs.jpcb.0c06477

