## Supplementary for "EnGens: a computational framework for generation and analysis of representative protein conformational ensembles"

#### Supplementary information

##### *FIGURES*

**Figure S1.** Structural view of PI3K-IA. The catalytic and regulatory subunits, as well as their respective subdomains, are presented below the structure.

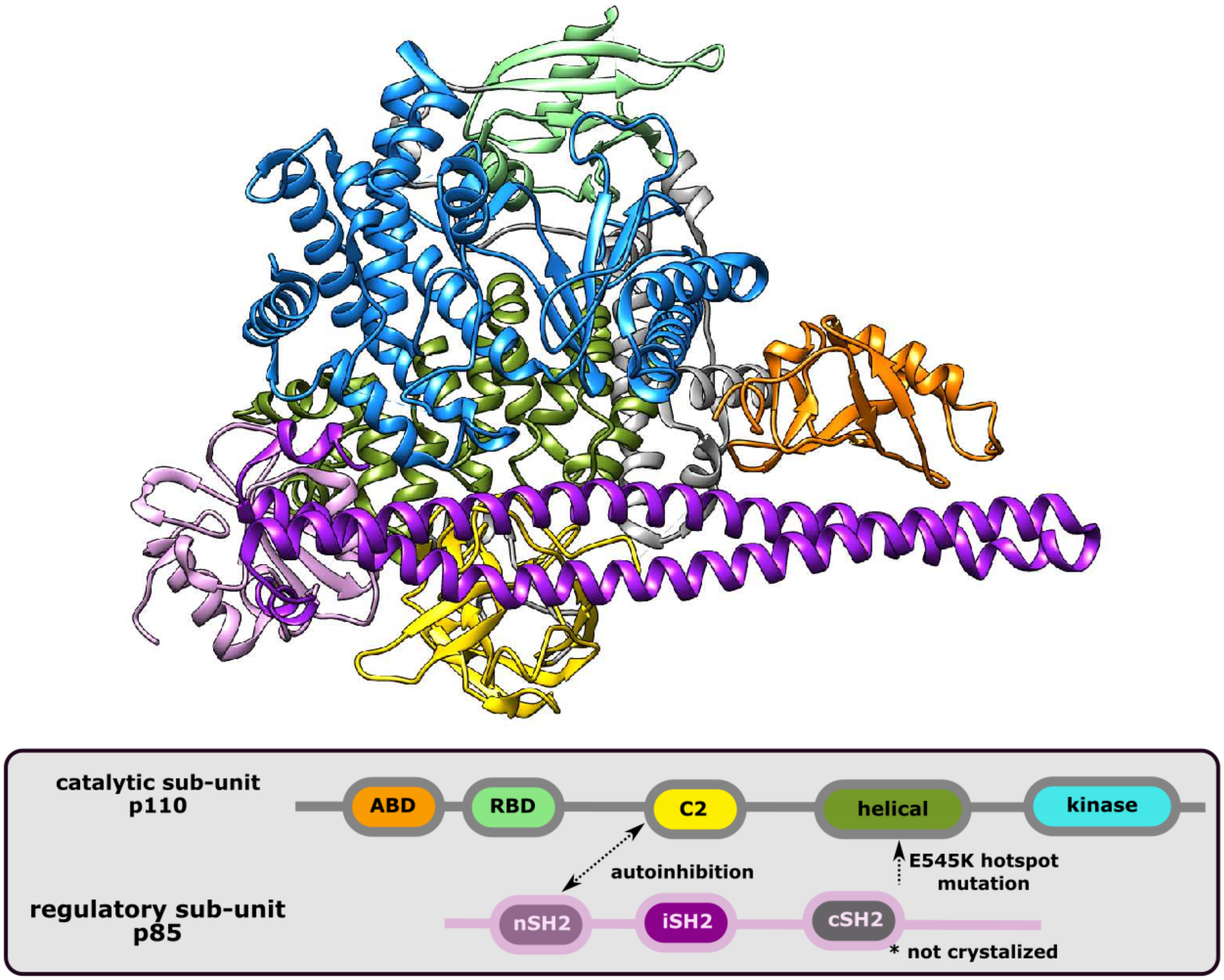

**Figure S2.** Result of the sequence mapping obtained from mTM-align multiple structure alignment for the PI3K regulatory subunit of the Zhang et al. dataset. Highlighted regions represent parts of the structures that do not contain gaps in any structure and that correspond to the extracted MCS.

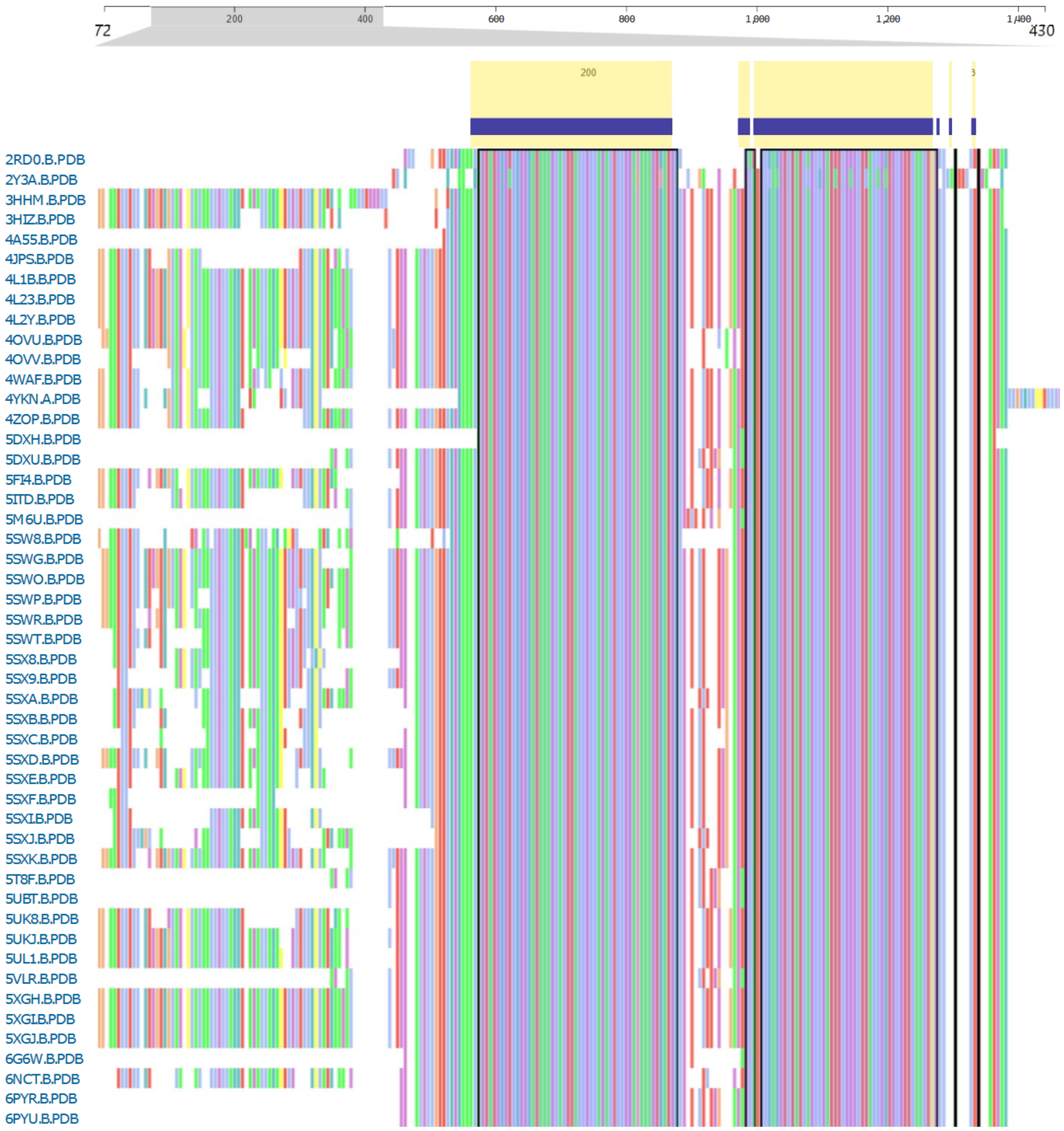

**Figure S3.** Result of the sequence mapping obtained from mTM-align multiple structure alignment for the PI3K catalytic subunit of the Zhang et al. dataset. Highlighted regions represent parts of the structures that do not contain gaps in any structure and that correspond to the extracted MCS.

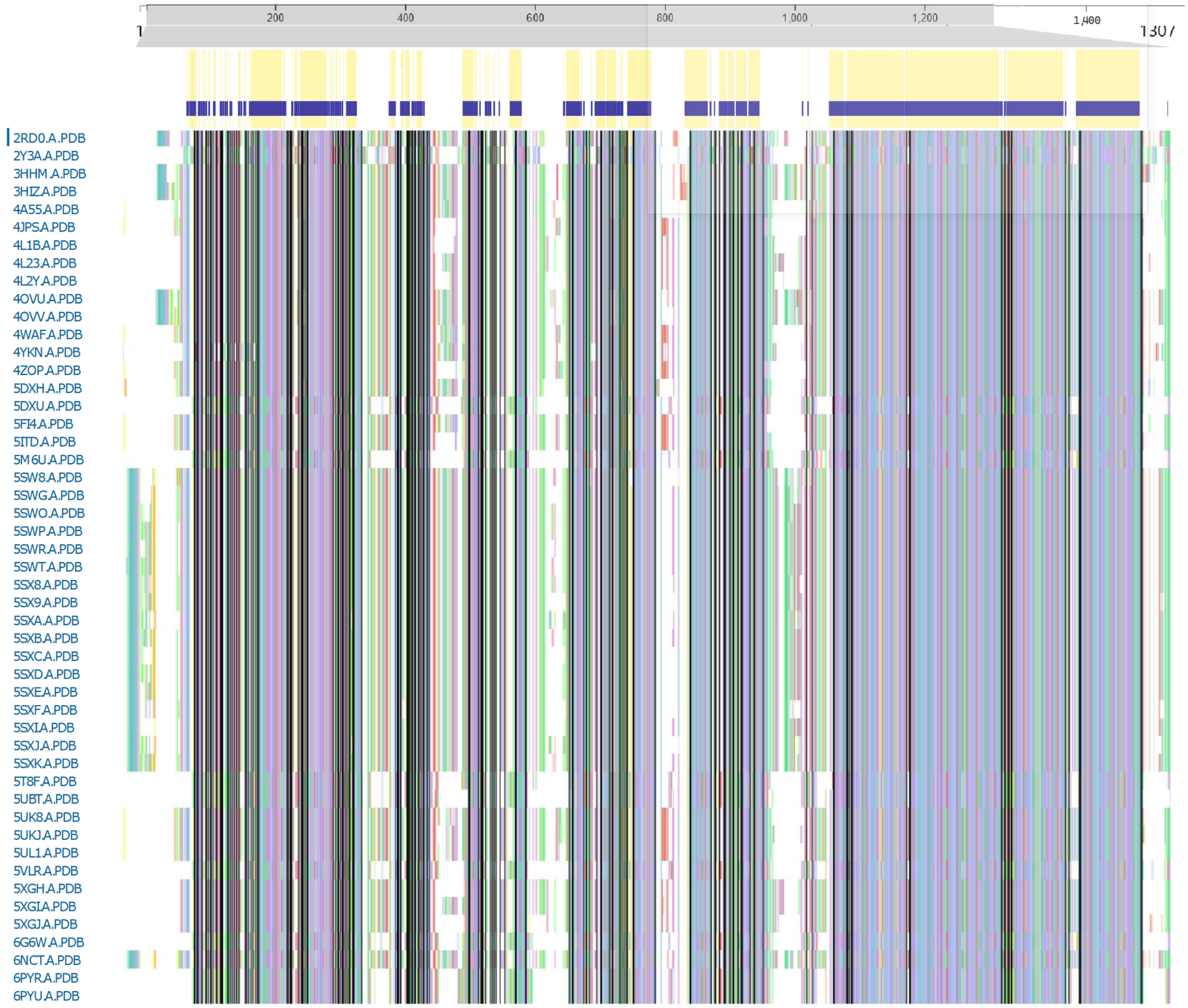

**Figure S4.** Original PI3K structure fetched from the PDB (pdb code: 4L1B). The MCS extracted based on the multiple structure alignment of the Zhang et al. dataset is highlighted in red.

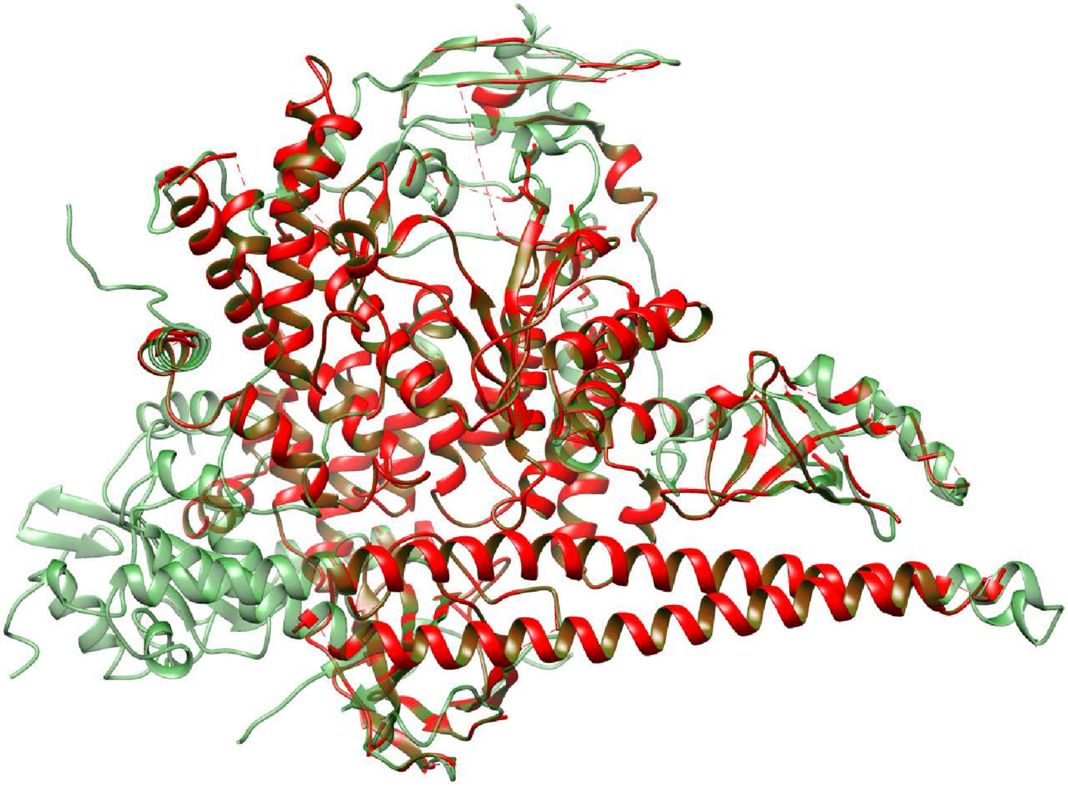

#### *TABLES*

**Table S1.** Comparison of the functionalities and availability of existing tools for generating and analyzing protein conformational ensembles. The tools compared to EnGens are CoNSEnsX, PENSA, ProDy.

| | *input data* | | *featurization* | | *dimensionality reduction* | | *availability* | | |
|---|---|---|---|---|---|---|---|---|---|
| *tool name* | *static* | *dynamic* | *predefined* | *custom* | *linear* | *nonlinear* | *Python package* | *Jupyter notebook* | *Docker image* |
| ***EnGens*** | ✔ | ✔ | ✔ | ✔ | ✔ | ✔ | ✔ | ✔ | ✔ |
| CoNSEnsX [1] | ✔ | ✖ | ✔ | ✖ | ✖ | ✖ | ✖ | ✖ | ✖ |
| PENSA [2] | ✖ | ✔ | ✔ | ✖ | ✔ | ✖ | ✔ | ✔ | ✖ |
| ProDy [3] | ✔ | ✔ | ✔ | ✖ | ✔ | ✖ | ✔ | ✖ | ✖ |

**Table S2.** PyEmma featurizations

| ***Feature name*** | ***Description*** | ***Required input arguments (PyEmma name)*** | ***Required input arguments (description)*** |
|---|---|---|---|
| all | Cartesian coordinates of all atoms | - | - |
| angles | Angles between custom selection of atoms | *indexes* | an array of triplets of atom indices between which the angles are evaluated |
| backbone torsions | Torsion angles of the backbone (phi and psi angles) | - | - |
| chi1 torsion | Chi1 angles | - | - |
| contacts | Contacts between atoms defined by the distance threshold of 0.3 Å | *indices* | an array of pairs of atoms between which contacts are calculated |

| | | | |
|---|---|---|---|
| | | | or<br><br>a list of atom indices (contacts are calculated between all pairs) |
| dihedrals | Dihedral angles between planes defined by a set of atoms | *indexes* | an array of quadruples of atoms for which dihedral angles are calculated |
| distances | Distances between pairs of atoms | *indices* | an array of pairs of atoms between which distances are calculated<br><br>or<br><br>a list of atom indices (distances are calculated between all pairs) |
| c-alpha distances | Distances between all c-alpha atoms | - | - |
| group COM | Centers of mass (COM) of groups of atoms | *group_definitions* | list of iterables representing groups of atoms |
| group mindist | Minimum distance between groups of atoms | group_definitions | list of iterables representing groups of atoms |
| inverse distances | Inverse distances between pairs of atoms | *indices* | an array of pairs of atoms between which distances are calculated<br><br>or<br><br>a list of atom indices (distances are calculated between all pairs) |

| | | | |
|---|---|---|---|
| minrmsd to reference | Minimum root mean square deviation (RMSD) to a reference structure | *ref* | reference structure |
| residue COM | Center of mass of residues | *residue_indices* | indices of residues |
| residue mindist | Minimum distance between pairs of residues (by default all pairs are used) | - | - |
| selection | Cartesian coordinates of the selection of atoms | *indexes* | indices of atoms included in the selection |
| sidechain torsions | Side-chain torsions | - | - |

**Table S3.** User inputs - hyper-parameters of the pipeline

| workflow | hyperparameter | description | default | required? |
|---|---|---|---|---|
| *WF 1S-2* | PDBFixer settings | Select how to clean the input structures. | add missing residues and atoms;<br><br>remove ligands, hydrogens, water, heteroatoms | no |
| *WF 1D-2* | align trajectory | Align all frames of the trajectory to a given structure. | aligning to the first frame of the trajectory | no (highly recommended) |
| *WF 1(S&D)-2* | selected region | Select a region of interest for analysis (e.g. binding site). All following actions will be performed on this selection. | no selection (whole structure is used) | no |

| | | | | |
|---|---|---|---|---|
| *WF 1(S&D)-3* | featurization | Select a way to embed the input structures(by choosing PyEmma featurizations from Table S2). | three default featurizations include:<br><br>- residue mindist<br>- backbone torsions<br>- residue mindist and backbone torsions | yes |
| *WF 1D-4* | VAMP-score featurization tuning | Decide if you want to tune the featurization based on VAMP-score. | The features providing the highest VAMP-score are selected from the default featurizations. | no |
| *WF 1D-4* | VAMP lag times | Select a range of lag times for calculating VAMP-score. | default: [0.5 ns, 1 ns, 2 ns]<br><br>recommended to set up your own lag times based on the length of the trajectory | yes (if VAMP featurization tuning is used) |
| *WF 1D-4* | VAMP dimensions | Select a number of components to be used for calculating VAMP-score | default: [2, 3, 4] | yes (if VAMP featurization tuning is used) |
| *WF 2(S&D)-2* | dimensionality reduction technique | Choose a techniques (PCA, UMAP, TICA or SRV). | default: UMAP | yes |
| *WF 2-2a* | variance threshold | Choose the threshold of the explained variance to select the top *k* PCs. | default: 95 | yes |

| | | | | |
|---|---|---|---|---|
| *WF 2-2b* | number of components | Choose the number of components onto which UMAP projects the data to. | default: 3 | yes |
| *WF 2D-2c* | lag time | Choose the set of lag times to perform TICA with. | default: a set of integers evenly spaced by 10, starting from 2 and ending at N/2 (N = length of the trajectory) | yes |
| *WF 2D-2c* | kinetic variance threshold | Choose the threshold of the explained kinetic variance to select the top k TICs. | default: 95 | yes |
| *WF 2D-2d* | lag time | Choose the set of lag times to perform SRV with. | default: a set of integers evenly spaced by 10, starting from 2 and ending at N/2 (N = length of the trajectory) | yes |
| *WF 2D-2d* | dimensionality | Choose the number of modes that SRV should find(equal to the number of processes) | default: 3 | yes (if the number of dimensions is insufficient, SRV might throw an error - in this case increase dimensionality) |
| *WF 3-2* | clustering technique | Choose one technique (K-means, GMM or hierarchical). | default: K-means | yes |
| *WF 3-2a* | centroid | Choose the | default: [2-10] | yes |

| | | | | |
|---|---|---|---|---|
| | counts | range for centroid counts - K-means will be performed for each value. | | |
| *WF 3-2a* | number of clusters | By inspecting the elbow plot or the silhouette score, pick the best number of clusters. | default: 3 | yes |
| *WF 3-2b* | component counts | Choose the range for component counts - GMM will be performed for each value. | default: [2-10] | yes |
| *WF 3-2b* | number of clusters | By inspecting the elbow plot or the silhouette score, pick the best number of clusters. | default: 3 | yes |
| *WF 3-2c* | linkage criterion | Choose the linkage criterion for agglomerative clustering. | default: ward | yes |
| *WF 3-2c* | number of clusters | By inspecting the dendogram, choose an appropriate number of | default: 3 | yes |

| | | | | |
|---|---|---|---|---|
| | | clusters. | | |
| *WF 3-3* | cluster weight threshold | Choose which clusters should be ignored by selecting a minimum weight | default: 0 (include all clusters) | yes |

**Table S4.** Zhang et al. dataset of dimer PI3K-IA crystal structures.

| Crystallized domains of the regulatory unit | Label | PI3K isoform | PDB codes (resolution) |
|---|---|---|---|
| nSH2, iSH2 | inactive/closed | alpha | 2rd0 (3.05 Å)<br>4jps (2.20 Å)<br>4a55 (3.50 Å)<br>4l1b (2.59 Å)<br>4l23 (2.50 Å)<br>4l2y (2.80 Å)<br>4ovu (2.96 Å)<br>4ovv (3.50 Å)<br>4waf (2.39 Å)<br>4ykn (2.90 Å)<br>4zop (2.62 Å)<br>5fi4 (2.50 Å)<br>5itd (3.02 Å)<br>5sw8 (3.30 Å)<br>5swg (3.11 Å)<br>5swo (3.50 Å)<br>5swp (3.41 Å)<br>5swr (3.31 Å)<br>5swt (3.49 Å)<br>5sx8 (3.47 Å)<br>5sx9 (3.52 Å)<br>5sxa (3.35 Å)<br>5sxb (3.30 Å)<br>5sxc (3.55 Å)<br>5sxd (3.50 Å)<br>5sxe (3.51 Å)<br>5sxf (3.46 Å)<br>5sxi (3.40 Å) |

| Crystallized domains of the regulatory unit | Label | PI3K isoform | PDB codes (resolution) |
|---|---|---|---|
| | | | 5sxj (3.42 Å)<br>5sxk (3.55 Å)<br>5uk8 (2.50 Å)<br>5ukj (2.80 Å)<br>5ul1 (3.00 Å)<br>5xgh (2.97 Å)<br>5xgi (2.56 Å)<br>5xgj (2.97 Å)<br>6nct (3.35 Å) |
| nSH2, iSH2 | active/open (through mutation) | alpha | 3hhm (2.80 Å)<br>3hiz (3.30 Å) |
| iSH2 | active/open | alpha | 5dxh (3.00 Å) |
| iSH2 | active/open | delta | 5dxu (2.64 Å)<br>5m6u (2.85 Å)<br>5t8f (2.91 Å)<br>5ubt (2.83 Å)<br>5vlr (2.80 Å)<br>6g6w (2.72 Å)<br>6pyr (2.21 Å)<br>6pyu (2.54 Å) |
| iSH2, cSH2 | active/open | beta | 2y3a (3.30 Å) |

### Workflows overview

- **Workflow 1S:** Extracting featurized representations from raw data (static use-case)
  - 1S-1: input description
  - 1S-2: preprocessing input
  - 1S-3: featurization
  - 1S-4: output description
- **Workflow 1D:** Extracting featurized representations from raw data (dynamic use-case)
  - 1D-1: input description
  - 1D-2: preprocessing input
  - 1D-3: featurization
  - 1D-4: VAMP featurization tuning
  - 1D-5: output description
- **Workflow 2(S&D)** Projecting the featurized representation into an embedding in low-dimensional space
  - 2-1: input description
  - 2-2: low-dimensionality reduction technique
    - 2-2a: PCA
    - 2-2b: UMAP
    - 2D-2c: TICA
    - 2D-2d: SRV
  - 2-3: output description
- **Workflow 3(S&D)** Clustering embeddings and extracting the ensemble
  - 3-1: input description
  - 3-2: clustering technique
    - 3-2a: K-Means
    - 3-2b: GMM
    - 3-2c: hierarchical
  - 3-3: representative selection
  - 3-4: output description
- **Workflow 4(S&D)** Visualizing the data and analyzing the ensemble
  - Embedding plot
  - Cluster membership plot
  - Multiple structures visualization
  - Ensemble summary box plots
  - Ensemble summary scatter plots

### Workflows detailed description

#### **Workflow 1S:** Extracting featurized representations from raw data (static use-case)

##### ***Workflow 1S-1:*** *input description*

The input for the first workflow in the static use-case can be given as any of the following:

- *List of files in the PDB format*
- *List of PDB codes* (corresponding PDB files are downloaded using pdb-tools package)
- *Uniprot IDs of a protein* (Protein Data Bank is queried for the PDB codes associated the protein with each given Uniprot ID using the PDB 1D Coordinate Server; the corresponding PDB files are fetched using pdb-tools package)

All of these inputs effectively result in a list of files in the PDB format, which contain Cartesian coordinates of all atoms in the protein structure. Before proceeding with the downstream workflows these input structures are preprocessed.

##### ***Workflow 1S-2:*** *preprocessing input*

Structures are first ***cleaned with the PDBFixer tool*** [4]. By default the preprocessing includes: adding missing residues and atoms; removing ligands, hydrogens, water and other heteroatoms. Users can edit these default settings.

Next, it is necessary to renumber the residues listed in these PDB files, so that each file involves the same numbering. ***Residues are renumbered using PDBrenum tool*** [5] according to their UniProt sequence.

Note that experimentally-resolved structures of the same protein may vary in the number of resolved residues. Some structures may contain additional chains and artifacts that helped stabilize the protein for crystallization. Others may lack regions that were not resolved. This is undesirable for downstream structural analysis. We want to analyze only the relevant parts of the structure and we want every structure to have the same number of residues. For these reasons, we first have to ***"trim" the structures to the same number of residues***. To this end, we perform multiple structure alignment of the input files (using mTM-align package [6]). As part of the structure alignment, mTM-align provides a sequence mapping where the amino acids of the sequences are matched to each other based on the structure alignment (e.g., Figures S2, S3) - we identify continuous regions of these amino acids (where no structure has a gap). These continuous regions form what we call the ***maximum common substructure*** (MCS). We extract the residues corresponding to the MCS from each structure (e.g., Figure S4). As a result, the input structures are *"trimmed"* to an MCS with a constant number of residues. Users can visually inspect this sequence mapping produced by the structure alignment and the extracted substructures (e.g., Figures S2, S3, S4) to make sure that they are valid.

Additionally, after extracting the MCS, users might be interested in analyzing only a specific part of a protein (e.g. only the binding pocket). We provide an option for ***selecting a region by identifying the residues of interest*** (using the selection algebra of the mdtraj package [7]). With this, the preprocessing of input structures is finalized and all the downstream analyses are performed on this preprocessed region of the protein.

***<u>Workflow 1S-3:</u>*** *<u>featurization</u>*

Each preprocessed input structure $s$ is still represented by the $x, y, z$ Cartesian coordinates of all its $N$ atoms - i.e., $s \in R^{3N}$. In other words, structure $s$ represents a point in configuration space $R^{3N}$. This representation is redundant - the motion of atoms is constrained by bonds and the underlying configuration space of the structure is actually a manifold of $R^{3N}$. Additionally, this representation is not invariant to translation and rotation.

Various mappings can be applied to embed each input structure into a different, more desirable space. Such mapping is often referred to as featurization, and can be viewed as:

$\chi: R^{3N} \rightarrow \Omega$ , where $\Omega$ is the configuration space of the featurized representation.

For example, $\chi$ can be applied to extract $\psi$ and $\varphi$ torsion angles of the protein backbone. In this case $\Omega$ has the non-Euclidian topology of a $r$-dimensional torus $T^r$ (where $r$ is the number of torsion angles). Note that standard algorithms for dimensionality reduction and clustering perform statistical analysis on Euclidean data, as Euclidean distance is used when calculating the mean and covariance for example. Approaches for dealing with non-Euclidean manifolds have been proposed, but are out of the scope of our current work. Thus, $\Omega$ must be Euclidean to comply with the downstream algorithms. A simple work-around recommended for dealing with angular values is to map angles to their sine and cosine values $\psi \rightarrow [sin(\psi), cos(\psi)]$. In this case, $\Omega$ would effectively be $R^{2r}$.

Mapping $\chi$ can be tailored to a specific protein and motivated by a biological insight. For example, it is known that the relative position of the nSH2 domain with respect to other domains of PI3K distinguishes active and inactive states of this enzyme. In this case an interesting featurization would contain pairwise distances between PI3K domains: $\chi$ can featurize the structure by calculating pairwise distances between the centers of mass of these domains. In that case $\Omega$ would correspond to $R^{\frac{d(d-1)}{2}}$ (where $d$ is the number of domains).

In both examples, the resulting featurized representations produced by $\chi$ will be invariant to rotations and translations of the structure in the original space.

We use the implementation of featurizations provided by the PyEmma package [8] that includes multiple featurization options summarized in Table S2.

***Workflow 1S-4:*** *output description*

After applying the featurization $\chi$ to each structure in the dataset, we combine all structures into a matrix $D_{n\times m}$, where $n$ is the number of input structures and $m$ is the dimensionality of the featurization. Note that $m$ is usually very high. For example, if the center of mass of the backbone C-alpha atoms is used for featurization of a protein with 300 residues, the resulting dimensionality $m$ will be 900. This motivates the second workflow, dimensionality reduction.

#### **Workflow 1D:** Extracting featurized representations from raw data (dynamic use-case)

***Workflow 1D-1:*** *input description*

In the dynamic use-case the input is the result of a molecular dynamics simulation, which includes the following:

- a trajectory (in any format accepted by PyEmma: .xtc, .dcd, .hdf5, .trr)
- a topology (in PDB format)

***Workflow 1D-2:*** *preprocessing input*

Frames in the trajectory should first be ***aligned*** (using mdtraj package [7]) to a reference frame. This is especially important when features dependent on translation and rotation (e.g., any featurization expressed in atomic Cartesian coordinates) are used, since small movements of the whole protein can add a lot of noise.

The next step involves ***selecting a region of interest*** (such as a binding site). Note that, contrary to the static use-case, here it is not necessary to trim the structures to an MCS - all the frames have exactly the same topology and the same number of residues.

***Workflow 1D-3:*** *featurization*

By default, each frame is represented by the Cartesian coordinates of all atoms. Similarly to the static use-case, featurization (see *Workflow 1S-3*) can be viewed as a transformation $\chi: R^{3N} \rightarrow \Omega$ applied to each frame. Frames can be better represented by any of the featurizations listed in Table S2.

***Workflow 1D-4:*** *VAMP featurization tuning*

Choosing the featurization is very important in both use-cases, as it can greatly impact the downstream analysis. In general, an optimal featurization will allow for identifying biologically-relevant and interesting states. One way to formalize this is to say that biologically relevant events are rare and that the processes that govern them are slow. A good featurization would allow for the identification of such slow processes.

Recent advances in Markov modeling of the system dynamics from MD data provide insights into resolving slow processes that guide the dynamics. Many approaches have been developed in an effort to model the dynamics from MD data, including time invariant component analysis (TICA), markov state modeling (MSM), variational approach to conformational dynamics (VAC), variational approach for learning Markov processes (VAMP). These methods are able to identify dominant (slow) dynamics by approximating the transfer operator guiding the dynamics. The VAMP approach approximates the Koopman transfer operator of the underlying Markovian dynamics by deriving its top singular components from the time series data. This approach is valuable because it can deal with both reversible and irreversible dynamics and because it provides VAMP-scores. The VAMP-E score quantifies the error in the approximated Koopman operator and can be used to tune the VAMP hyper-parameters. One of these hyper-parameters is the featurization. Thus, the featurization that yields the maximum VAMP-E score allows for the most accurate resolution of the dynamics and can be viewed as optimal.

Here, we ***use PyEmma's implementation of VAMP scores*** to allow users to evaluate their featurization by calculating the VAMP-E score. Users can ***choose amongst a range of different featurizations by picking the one with the highest VAMP score***.

***Workflow 1D-5:*** *output description*

After applying the selected featurization to each frame in the trajectory - frames are combined into a matrix $D_{nxm}$ , where $n$ is the number of frames and $m$ is the dimensionality of the featurization. This matrix is similar to the matrix resulting from Workflow 1S and almost all downstream analyses will be the same for the static use-case (S) and the dynamic use-case (D). The only exceptions are dimensionality reduction techniques in steps 2D-2c and 2D-2d (i.e., TICA and SRV). These methods model the dynamics of the data and can thus be applied only to the dynamic use-case.

#### **Workflow 2(S&D):** Projecting the featurized representation into an embedding in low-dimensional space (static and dynamic use-case)

***Workflow 2-1:*** *input description*

The input to Workflow 2 consists of the featurized dataset expressed as the matrix $D_{n\times m}$. For the static use-case $n$ corresponds to the number of collected structures, for the dynamic use-case $n$ is the number of frames in the trajectory. $m$ corresponds to the dimensionality of the chosen featurization. Since $m$ is usually high as the featurizations have high dimensionality, the important step executed in Workflow 2 is reducing this dimensionality by projecting the data into a lower dimensional space by preserving important properties of the data expressed as the variance or the kinetic variance.

Different methods offered by EnGens are presented in following subsections including PCA (WF2-2a), TICA (WF2-2b), UMAP (WF2D-2c) and SRV (WF2D-2d). Note that PCA and TICA are linear methods that find a linear projection into a lower-dimensional space, while UMAP and SRV perform a non-linear projection. In addition, TICA and SRV use the dynamic properties of the underlying MD datasets and should not be applied to the static use-case.

***Workflow 2-2a:*** *PCA*

Principal component analysis (PCA) is a widely used and powerful statistical technique for reducing dimensionality. The key idea of PCA is in finding the "principal components" (PCs - vectors in high-dimensional spaces) onto which the data can be projected while preserving as much variance as possible.

The first step is to normalize the data by subtracting the mean and the standard deviation of the dataset from each point. This is important to ensure that individual feature variances are comparable. Next, the eigenvectors of the data metrics are calculated using singular value decomposition methods. These eigenvectors correspond to PCs. Next, the data is projected onto the PCs. The *k* PCs with the highest eigenvalues are selected, effectively reducing the dimensionality of the input data to *k* dimensions. Usually, *k* is chosen such that a satisfactory amount of variance is retained (e.g., 90%).

EnGens uses scikit-learn implementation of the PCA method to obtain the final matrix $D_{n\times k}$ where n is the number of structures (or frames) in the dataset and k is the number of chosen PCs. Note that PCA is a linear method: it assumes that the relationship between features can be represented by a linear combination of the original features. If a nonlinear relationship between features is suspected, a more suitable method for dimensionality reduction might be UMAP.

***Workflow 2-2b:*** *UMAP*

The Uniform Manifold Approximation and Projection (UMAP) algorithm [9] is a non-linear dimensionality reduction technique that can preserve the global structure of the data. UMAP aims to approximate the underlying manifold of the data by constructing a fuzzy topological representation of these data and projecting them onto a lower-dimensional space.

The first step in UMAP is the construction of the weighted fuzzy graph. This step is controlled by parameters: n_neighbors and min_dist. These parameters indicate the number of nearest neighbors that each point is connected to and the minimum distance between neighbors. Next, the spectral embedding of the constructed graph is calculated and optimized with respect to the fuzzy set cross-entropy (or its differentiable approximation) using stochastic gradient descent. The terms of the fuzzy set cross-entropy objective function can be viewed as attractive and repulsive forces. Optimizing this objective will balance the "pull" and "push" of the data and the low-dimensional representation should settle into a state that relatively accurately represents the overall topology of the original data. The n_components parameter indicates the dimensionality of the embedding and it has to be selected prior to fitting UMAP.

EnGens uses an open-source UMAP implementation [9] to produce the final matrix $D_{n\times k}$ where n is the number of structures (or frames) in the dataset and k is equal to n_components (the number of chosen components). Note that UMAP assumes that the data are uniformly distributed on the underlying manifold. Consequently, it does not preserve the relative local density of the data well. This fact is highlighted when the two-dimensional free energy plots of UMAP projections are generated, as they rely on estimating the density of clusters. Thus, when using UMAP for dimensionality reduction, all downstream analyses of density estimation should be performed with care.

***Workflow 2D-2c:*** *TICA*

Time-invariant component analysis (TICA) [10] is a dimensionality reduction technique commonly used in the analysis of time series data (such as MD trajectory data). Similarly to PCA, TICA finds the components (vectors) onto which all datapoints should be projected. However, the objective of TICA is not to find the components preserving most of the variance. Instead, TICA aims to identify slow dynamics that govern the behavior of the system. To this end, TICA looks for a coordinate system by identifying the linear combinations of the original coordinates that change most slowly over time.

The optimization problem formalized by TICA is to find coordinates of maximal autocorrelation at the given lag time. This can be achieved by the following few steps. First, the mean-free covariance ($C_0$) and mean-free time-lagged covariance ($C_\tau$) matrices are calculated from the data. Next, the generalized eigenvalue problem is solved $C_\tau r_i = C_0 \lambda_i r_i$, where $r_i$ are the eigenvectors corresponding to time independent components - TICs, while $\lambda_i$ are the respective eigenvalues that indicate the relaxation time-scales of the TICs. Finally, the k slowest resolved processes (TICs) are picked to project the data. Choosing $k$ can be done by setting a threshold on the kinetic variance that should be retained in the data.

EnGens uses PyEmma's implementation of TICA to produce the final matrix $D_{n\times k}$ where n is the number of frames in the trajectory and $k$ is the number of chosen TICs. Note that TICA is a linear method and assumes a linear relationship between features. For resolving non-linear slow processes the SRV method is recommended.

***Workflow 2D-2d:*** *SRV*

The state-free reversible VAMPnet (SRV) [11] is a non-linear dimensionality reduction technique. Similarly to TICA, SRV aims to identify the slow processes (modes) that drive the dynamics of the system. In contrast to TICA, which solves a generalized eigenvalue problem, SRV resolves the processes by *learning* the leading eigenfunctions of a dynamical system transfer operator from trajectory data. Using a deep learning architecture allows SRV to identify non-linear processes.

In SRV, a Siamese neural network is constructed with two subnets that share the same architecture and weights. A pair of datapoints is fed into the network. The pairs are selected to be rows of the featurized matrix $D_{n\times m}$separated by the lag time ($d_t$, $d_{t+\tau:}$). The network learns the mappings $f_i$, where $i = (1,.., k)$ and $k$ is the number of modes we want to learn. This parameter is set by the user; it defines the architecture as the number of nodes in the last hidden layer of the network will be $k$. The network produces the outputs ($f_i(d_t)$, $f_i(d_{t+\tau:})$). The loss function is calculated using these outputs and it corresponds to the negative VAMP-2 score. As the network maximizes the VAMP-2 score, it learns non-linear feature embeddings that allow for the best approximation of the transfer operator. In effect, this results in learning a good non-linear low-dimensional embedding of the original features.

EnGens uses the open-source implementation of SRV to produce the final matrix $D_{n\times k}$ where n is the number of frames in the trajectory and $k$ is the selected number of modes. When SRV is used, the embedding dimensionality $k$ and the lag time τ have to be carefully selected by the user.

***Workflow 2-3:*** *output description*

After projecting the featurized representation of each datapoint to a lower-dimensional embedding, the resulting data matrix has the form $D_{nxk}$ , where $n$ is the number of input structures or the number of frames in the trajectory, and $k$ is the dimensionality of the featurization.

#### **Workflow 3(S&D):** Clustering embeddings and extracting the ensemble (static and dynamic use-case)

***Workflow 3-1:*** *input description*

For both the dynamic and static use-case, the input to Workflow 3 consists of the embedded dataset expressed as the matrix $D_{n\times k}$. For the static use-case $n$ corresponds to the number of collected structures, for the dynamic use-case $n$ is the number of frames in the trajectory. $k$ corresponds to the dimensionality of the embedding. After reducing the dimensionality of the data in Workflow 2 it is possible to visualize the dataset in a 2D or 3D space defined by the top identified component, which gives better insight into the dataset. The additional step of clustering performed in Workflow 3 allows for the data to be stratified into groups of datapoints (structures) that are close together in this embedded space and are assumed to be structurally similar. To this end, users can choose one of these widely used methods: K-means, hierarchical or GMM clustering.

***Workflow 3-2a:*** *K-means clustering*

The K-means algorithm aims to separate $n$ datapoints into $c$ clusters by assigning each datapoint to one of the $c$ centroids. First, the positions of the centroids are initialized (randomly or via some heuristic). Next, the algorithm iterates between: (1) generating clusters by assigning points to the closest centroid, and (2) repositioning the centroids to the mean of the clusters. The centroid assignment is evaluated using inertia (or sum of distances within a cluster): $\sum_{i=0}^{n} min_{\mu_j \in C} || d_i - \mu_j ||$, where $i$ iterates through all $n$ datapoints ($d_i$) and $\mu_j$ is the centroid (from the set $C$ of all centroids) to which $d_i$ is assigned. The algorithm iterates until the movement of the centroids between subsequent steps is under some threshold (i.e., until convergence).

The number of centroids $c$ (i.e., the number of produced clusters) has to be predefined. This value is often difficult to predict and it is recommended users evaluate different values. After running K-means with a range of $c$ values, the best $c$ can be selected by plotting the corresponding inertia and using the elbow method (finding the $c$ for which inertia drops rapidly). Using the elbow method, $c$ will be selected such that clusters are compact around the centroids. Alternatively, users can inspect the silhouette plots for different $c$ values and select the value with the maximum average silhouette score. The silhouette score is assigned to each datapoint $i$ as $s(i) = \frac{b(i)-a(i)}{max\{a(i), b(i)\}}$, where $b(i)$ is the mean distance between point $i$ and points from the nearest cluster and $a(i)$ is the mean distance between point $i$ and points from its own cluster. The silhouette score is in the range (-1, +1) where -1 indicates overlapped clusters and +1 indicates tight, well separated clusters. Maximizing the silhouette score will result in picking the clustering with clear separation between datapoints and little overlap between clusters.

EnGens uses scikit-learn [12] implementation of the K-means algorithm to generate the cluster assignment of all datapoints. Note that K-means uses inertia to evaluate clusters, which relies on convex and isotropic assumptions. This results in spherical clusters, which might not always be a good fit for the data. Alternatively, if the underlying data distribution is Gaussian, GMM clustering can be used to produce clusters of different shapes, sizes, and orientations.

***Workflow 3-2b:*** *GMM clustering*

Gaussian Mixture Models are probabilistic generative models that can represent a dataset as a mixture of Gaussian distributions with unknown parameters. The number of distributions (components) is predefined (similarly to the number of centroids in K-means). The expectation-maximization (EM) algorithm is employed to fit the parameters of the Gaussian distributions to the data. EM consists of consecutive steps of (E) calculating the log-likelihood of the data given the current set of parameters and (M) finding new parameters that maximize the given log-likelihood. After modeling the data as a mixture of Gaussian distributions, each datapoint ends up associated with a probability for each of the GMM components. Eventually, datapoints are assigned to the GMM component (i.e., the cluster) with the highest associated probability.

The number of GMM components is predefined. Users can optimize this number by running GMM on a range of values and picking the one with the lowest associated Bayesian information criterion (BIC). BIC is defined as $k\ ln(N) - 2ln(\widehat{L})$, where $k$ is the number of GMM components, $N$ is the number of datapoints and $\widehat{L}$ is the likelihood of the data under the inferred GMM model. BIC penalizes model complexity with the first term (preventing overfitting) and rewards model fitting with the second term. Analysis of BIC allows picking k using the elbow method (finding the number of components for which BIC drops rapidly). Alternatively, clustering can be assessed with the silhouette method in a similar manner as for the K-means method.

EnGens uses scikit-learn [12] implementation of the GMM algorithm to produce the cluster assignment of the datapoints. Note that both K-means and GMM require careful consideration of the number of resulting clusters, and that (a range of values for) this parameter has to be defined before running the algorithm. On the other hand, hierarchical clustering uses a different approach which allows defining the number of clusters after computing a dendrogram of the data.

***Workflow 3-2c:*** *hierarchical clustering*

Hierarchical clustering forms groups of datapoints by successively merging or dividing the dataset. In EnGens, we employ the agglomerative hierarchical clustering, which is a bottom-up approach which iteratively merges datapoints into groups. First, each datapoint defines its own cluster. Next, clusters are iteratively merged based on a given criterion. For example, when the *average linkage* criterion is used, a pair of clusters is merged if the average distances between all their points are minimum. Merging steps are performed until only one cluster remains. The resulting dendrogram can be inspected to choose the number of clusters that best describes the data.

EnGens uses scikit-learn [12] implementation of the agglomerative hierarchical clustering. Note that for a large number of datapoints, agglomerative clustering can be computationally intensive, which might be of particular concern for MD data with a high number of frames. To combat this issue, users can provide a subsample of the trajectory as input (e.g., by loading every 10th frame).

***Workflow 3-3:*** *representative selection*

After clustering, the input matrix $D_{n\times k}$, each cluster contains a set of datapoints (structures) that are assumed to be structurally similar. In Workflow 3-3, a single datapoint, called the cluster representative, is selected from each cluster to represent it within the final representative ensemble. If users do not want to pick representatives from all clusters (e.g., to ignore very small clusters), a weight threshold can be used to discard clusters.

There are two ways to pick a representative from each cluster. First, the representative can be the point closest to the center of the cluster: in K-means, cluster centroids are the centers; in GMM, the means of the distributions are the centers.. Alternatively, the representative can be a *hub* of the cluster, which is defined as the point with the most neighbors in the cluster. Neighbors are defined as datapoints that are closer to the hub than the mean of pairwise distances within the cluster.

***Workflow 3-3:*** *output description*

The result of this workflow is (1) the grouping of the datapoints into clusters of structures with similar properties and (2) the final representative ensemble composed of the extracted cluster representatives.

#### **Workflow 4(S&D):** Visualizing the data and analyzing the ensemble

EnGens provides a variety of plots to summarize the resulting data clustering and generated representative ensemble. Plots are implemented using Plotly and NGLViewer[13] is used to visualize the 3D structures of the representatives.

***Embedding plot***

This plot shows the projected embedding from Workflow 2 in two dimensions. The X-axis represents the first principal component (or TIC/SRV/UMAP component) while the Y-axis represents the second principal component (or TIC/SRV/UMAP component). Each point corresponds to one structure (or frame) from the dataset. Points are colored based on their cluster membership.

***Cluster membership plot***

This plot shows the details of cluster membership across the whole dataset. Y-axis represents cluster membership. In the static case, the X-axis lists the structure IDs in the order specified in the original dataset. In the dynamic case, the X-axis lists the MD frames according to the simulation time.

***Multiple structures visualization***

We use NGLViewer to show the extracted representative ensemble within the notebook. It is possible to select a cluster of interest and display the 3D cartoon representation of the cluster representative. In addition, other uniformly sampled structures from the same cluster can be visualized.

***Ensemble summary box plots***

This plot summarizes a property of interest (e.g., RMSD to a reference structure) across different clusters with a box plot. The X-axis lists the cluster IDs. The Y-axis corresponds to the selected property.

***Ensemble summary scatter plots***

This plot summarizes a property of interest (e.g., RMSD to a reference structure) across different clusters with a scatterplot. The X-axis lists the structure IDs (i.e, the number of an MD frame in the dynamic data, or a structure name in the static data). The Y-axis corresponds to the selected property. Points are colored based on their cluster membership. Selected representative structures are highlighted in red.

### References


[1] A. F. Ángyán, B. Szappanos, A. Perczel, and Z. Gáspári, "CoNSEnsX: an ensemble view of protein structures and NMR-derived experimental data," *BMC Structural Biology*, vol. 10, no. 1, p. 39, Oct. 2010, doi: 10.1186/1472-6807-10-39.

[2] M. Vögele, N. J. Thomson, S. T. Truong, J. McAvity, U. Zachariae, and R. O. Dror, "Systematic Analysis of Biomolecular Conformational Ensembles with PENSA." arXiv, Dec. 05, 2022. doi: 10.48550/arXiv.2212.02714.

[3] S. Zhang *et al.*, "ProDy 2.0: increased scale and scope after 10 years of protein dynamics modelling with Python," *Bioinformatics*, vol. 37, no. 20, pp. 3657–3659, Oct. 2021, doi: 10.1093/bioinformatics/btab187.

[4] P. Eastman *et al.*, “OpenMM 7: Rapid development of high performance algorithms for molecular dynamics,” *PLoS Comput Biol*, vol. 13, no. 7, p. e1005659, Jul. 2017, doi: 10.1371/journal.pcbi.1005659.

[5] B. Faezov and R. L. D. Jr, “PDBrenum: A webserver and program providing Protein Data Bank files renumbered according to their UniProt sequences,” *PLOS ONE*, vol. 16, no. 7, p. e0253411, Jul. 2021, doi: 10.1371/journal.pone.0253411.

[6] R. Dong, Z. Peng, Y. Zhang, and J. Yang, “mTM-align: an algorithm for fast and accurate multiple protein structure alignment,” *Bioinformatics*, vol. 34, no. 10, pp. 1719–1725, May 2018, doi: 10.1093/bioinformatics/btx828.

[7] R. T. McGibbon *et al.*, “MDTraj: A Modern Open Library for the Analysis of Molecular Dynamics Trajectories,” *Biophys J*, vol. 109, no. 8, pp. 1528–1532, Oct. 2015, doi: 10.1016/j.bpj.2015.08.015.

[8] M. K. Scherer *et al.*, “PyEMMA 2: A Software Package for Estimation, Validation, and Analysis of Markov Models,” *J. Chem. Theory Comput.*, vol. 11, no. 11, pp. 5525–5542, Nov. 2015, doi: 10.1021/acs.jctc.5b00743.

[9] L. McInnes, J. Healy, N. Saul, and L. Großberger, “UMAP: Uniform Manifold Approximation and Projection,” *Journal of Open Source Software*, vol. 3, no. 29, p. 861, Sep. 2018, doi: 10.21105/joss.00861.

[10] C. R. Schwantes and V. S. Pande, “Modeling Molecular Kinetics with tICA and the Kernel Trick,” *J. Chem. Theory Comput.*, vol. 11, no. 2, pp. 600–608, Feb. 2015, doi: 10.1021/ct5007357.

[11] W. Chen, H. Sidky, and A. L. Ferguson, “Nonlinear discovery of slow molecular modes using state-free reversible VAMPnets,” *J. Chem. Phys.*, vol. 150, no. 21, p. 214114, Jun. 2019, doi: 10.1063/1.5092521.

[12] F. Pedregosa *et al.*, “Scikit-learn: Machine Learning in Python,” *Journal of Machine Learning Research*, vol. 12, no. 85, pp. 2825–2830, 2011.

[13] A. S. Rose, A. R. Bradley, Y. Valasatava, J. M. Duarte, A. Prlić, and P. W. Rose, “NGL viewer: web-based molecular graphics for large complexes,” *Bioinformatics*, vol. 34, no. 21, pp. 3755–3758, Nov. 2018, doi: 10.1093/bioinformatics/bty419.
